# Spatial specificity of the functional MRI blood oxygenation response relative to neuronal activity

**DOI:** 10.1101/097287

**Authors:** Denis Chaimow, Essa Yacoub, Kâmil Uğurbil, Amir Shmuel

## Abstract

Previous attempts at characterizing the spatial specificity of the blood oxygenation level dependent functional MRI (BOLD fMRI) response by estimating its point-spread function (PSF) have conventionally relied on spatial representations of visual stimuli in area V1. Consequently, their estimates were confounded by the width and scatter of receptive fields of V1 neurons. Here, we circumvent these limits by instead using the inherent cortical spatial organization of ocular dominance columns (ODCs) to determine the PSF for both Gradient Echo (GE) and Spin Echo (SE) BOLD imaging at 7 Tesla. By applying Markov Chain Monte Carlo sampling on a probabilistic generative model of imaging ODCs, we quantified the PSFs that best predict the spatial structure and magnitude of differential ODCs’ responses. Prior distributions for the ODC model parameters were determined by analyzing published data of cytochrome oxidase patterns from post-mortem histology of human V1 and of neurophysiological ocular dominance indices. The most probable PSF full-widths at half-maximum were 0.82 mm (SE) and 1.02 mm (GE). Our results provide a quantitative basis for the spatial specificity of BOLD fMRI at ultra-high fields, which can be used for planning and interpretation of high-resolution differential fMRI of fine-scale cortical organizations.

## Introduction

Functional magnetic resonance imaging (fMRI) of the human brain is increasingly being used to investigate fine-scale structures such as cortical columns (Cheng et al., 2001; De Martino et al., 2015; Goodyear and Menon, 2001; Menon et al., 1997; Shmuel et al., 2010; Yacoub et al., 2008; 2007; Zimmermann et al., 2011). To optimally plan high-resolution fMRI studies and to correctly interpret their results it is necessary to know the inherent limits of the fMRI spatial specificity relative to the sites where changes in neuronal activity occur.

The most commonly used fMRI approach relies on gradient echo (GE) blood oxygenation level dependent (BOLD) contrast (Bandettini et al., 1992; Kwong et al., 1992; Ogawa et al., 1990; 1992). GE BOLD is sensitive to the intra- and extravascular effects of activation-induced changes in the deoxyhemoglobin content of blood. At standard magnetic field strengths (1.5 T, 3 T) the signal is dominated by contributions from larger blood vessels. At higher magnetic field strengths the strong intravascular component of these large blood vessels decreases, while the extravascular signal changes around capillaries and smaller vessels increase ( Uludağ et al., 2009; Yacoub et al., 2001). Additional weighting towards the microvasculature can be achieved by using spin echo (SE) BOLD imaging, which suppresses extravascular signal contributions from larger blood vessels ( Uludağ et al., 2009; Yacoub et al., 2003).

The first study to quantify the spatial specificity of the BOLD response (Engel et al., 1997) used an elegant phase-encoding paradigm that induced traveling waves of retinotopic neural activity in the primary visual cortex (V1). Assuming a shift-invariant linear response, Engel et al. (1997) estimated the point-spread function (PSF), which represents the spatial response that would be elicited by a small point stimulus. They found the full-width at half-maximum (FWHM) of the GE BOLD PSF to be 3.5 mm at 1.5 T. Similar values (3.9 mm for GE BOLD and 3.4 mm for SE BOLD) have been reported at 3 T (Parkes et al., 2005) using a paradigm similar to that used in Engel et al. (1997). To estimate the GE BOLD PSF at 7 T, we previously measured the spatiotemporal spread of the fMRI response in grey matter regions around the V1 representation of edges of visual stimuli (Shmuel et al., 2007). To reduce contributions from macroscopic veins, we excluded voxels that showed vessel-like response features. The mean measured and estimated FWHMs were 2.34 ± 0.20 mm and < 2 mm, respectively. The spatial specificity of SE BOLD fMRI at ultra-high magnetic fields has not yet been quantified.

All previous attempts at characterizing the spatial specificity of the BOLD fMRI response (Engel et al., 1997; Parkes et al., 2005; Shmuel et al., 2007) relied on an implicit assumption that neuronal responses to small visual stimuli are point-like. However, to estimate the spatial specificity of the BOLD response, these studies have conventionally relied on spatial representations of visual stimuli in area V1. Unlike the implicit assumption of point-like responses, the receptive fields of neurons in V1 have non-zero spatial extents (Hubel and Wiesel, 1968). In addition, electrode measurements in macaque V1, oriented orthogonally relative to the surface of cortex have demonstrated substantial scatter in the center of receptive fields (Hubel and Wiesel, 1974). Therefore, the pattern of neural activity parallel to the cortical surface is a blurred representation of the visual stimulus. This implies that receptive field size and scatter pose a lower limit on any BOLD fMRI PSF width that is estimated using spatial representations of visual stimuli in V1. Consequently, the previously computed estimates of the spatial specificity of the fMRI response were confounded by the width and scatter of receptive fields of V1 neurons. Such estimates are limited in that they solely measure the capacity of the BOLD response to resolve retinotopic representations; they do not measure its ability to resolve more fine-grained neural activity. Yet only this latter resolvability matters for functional imaging at the spatial scale of cortical columns.

Here, we estimate and compare the PSF widths of GE and SE BOLD imaging at 7 T using a novel approach. We circumvent the limits posed by the retinotopic representation of visual stimuli by instead using the inherent cortical spatial organization of ocular dominance columns (ODCs). To this end, we fit a model of ODCs imaging (Chaimow et al., 2011) to ODCs responses acquired at 7 T (Yacoub et al., 2007). We quantify the width of the PSF that best predicts the spatial structure and magnitude of differential ODC responses. Since we do not have access to the underlying anatomical ODC patterns and neurophysiological responses, we use a probabilistic modeling approach. We constrain the model ODC parameters by estimating features of real ODC patterns taken from post-mortem cytochrome oxidase (CO) maps of human ODCs (Adams et al., 2007) and neurophysiological response distributions in primates (Berens et al., 2008; Hubel and Wiesel, 1968). We then fit our model by Markov Chain Monte Carlo (MCMC) sampling. Our results provide a quantitative basis for the spatial specificity of differential BOLD fMRI at ultra-high fields.

## Methods

### Overview

We developed a probabilistic generative model of imaging ODCs in order to estimate widths of GE and SE BOLD PSFs that would best explain our previously obtained fMRI data of differential ODC maps (Yacoub et al., 2007).

The measured fMRI maps consisted of voxel estimates of the difference in BOLD responses to left and right eye stimulation. We modeled these responses as the sum of predictions from an ODC imaging model and measurement noise. The predictions from the ODC imaging model were completely determined by a set of parameters. Accounting for the effect of the measurement noise allowed us to first express the probability of observing the measured fMRI maps as a function of model parameters. In a second stage, we derived the posterior probability of the model parameters given the observed data and the prior probability of parameters.

### Model of imaging ODCs

We implemented a model of imaging ODCs (Chaimow et al., 2011; see Fig. 1 for an overview; see Appendix A in the Suppl. Material for the detailed equations). The first component of the model, i.e. the modeling of realistic ODCs, followed (Rojer and Schwartz, 1990). It consisted of band-pass filtering of spatial white noise using an anisotropic filter. The filtering was followed by applying a sigmoidal point-wise non-linearity, which controlled the smoothness of transitions between left and right eye preference regions.

**Fig. 1.**
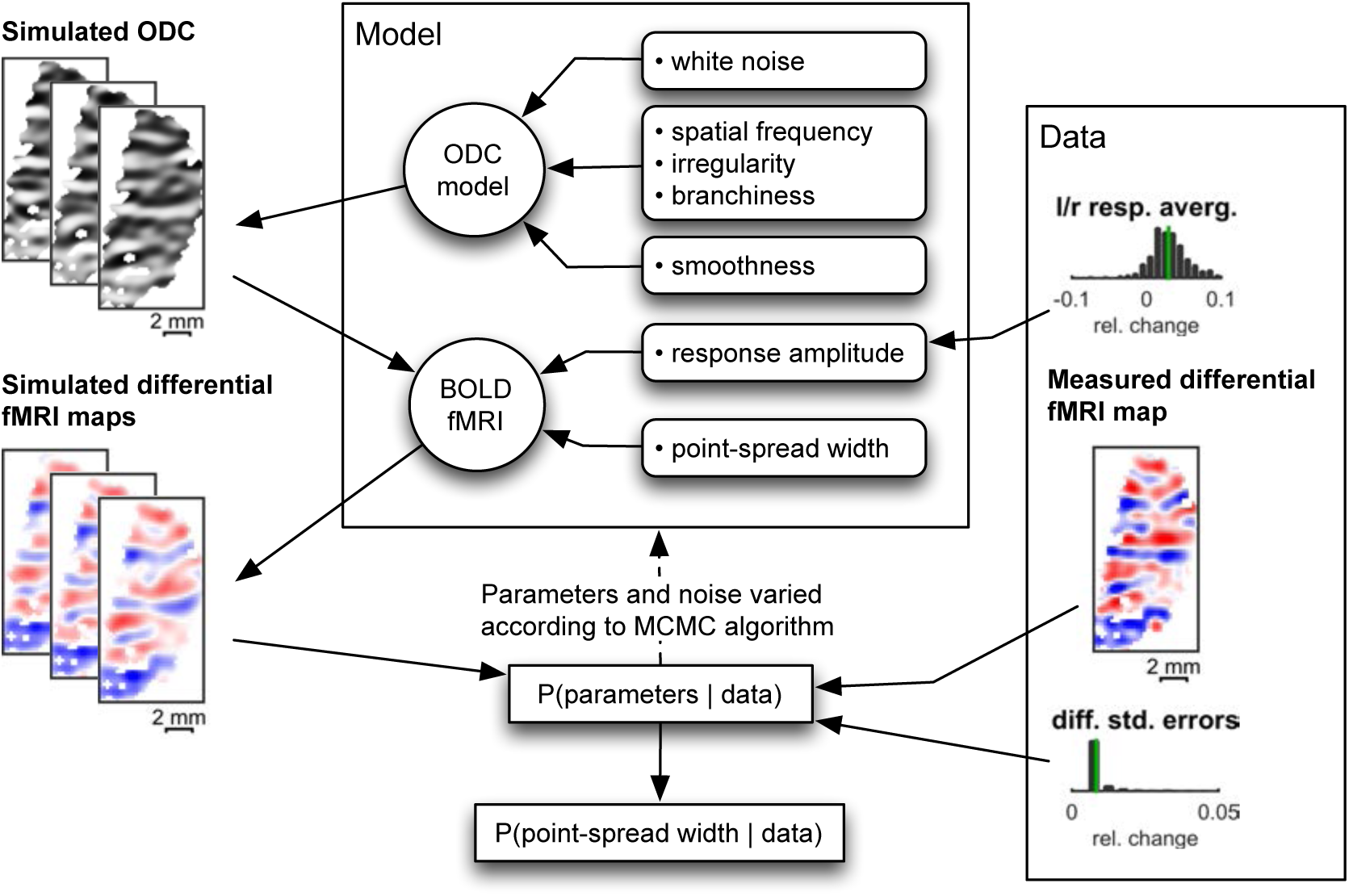
Overview of Markov Chain Monte Carlo fitting. The model was fitted to the fMRI data using Markov Chain Monte Carlo (MCMC) sampling. For an arbitrary given set of parameters, the model generated a differential fMRI map (left). This map was compared to the measured fMRI map (right) and the likelihood of parameters given the data was calculated. The MCMC algorithm uses this likelihood together with parameter priors to further traverse the parameter space. After sufficiently many iterations the resulting parameter samples are distributed according to their joint posterior probability distribution.

The spatial BOLD response was modeled as a convolution of the ODCs pattern with a Gaussian PSF. We modeled separate BOLD responses for the GE and SE maps. However, as these maps were obtained from the exact same region in area V1 in each subject, the model constituted one ODC map underlying both the GE and SE responses. MRI k-space sampling was modeled by restricting the spatial frequency space to its central part in accordance with the modeled field of view, sampling matrix and voxel size (Haacke et al., 1999).

All model parameters are listed in Table 1. Three parameters controlled general ODC properties: the main (peak) spatial frequency ρ, the degree of irregularity δ and the branchiness ϵ. The spatial white noise served as a high-dimensional parameter determining the specific manifestation of the ODC pattern. A parameter *ω* controlled the smoothness of transitions between regions showing left and right eye preference. The PSFs of the BOLD responses were parameterized by amplitudes *𝛽_GE_* and *𝛽_SE_* and by their FWHM *fwhm_GE_* and *fwhm_SE_* (the parameters of interest).

Relative to our previously published model, we made 2 slight modifications. First, to simplify derivation of the gradient we subsequently used for implementing the MCMC sampling, we modified the formulation of the bandpass filtering kernel (see Appendix A). Second, due to consideration of step size determined by the MCMC algorithm, we defined and used a smoothness parameter *ω* instead of using its inverse, the sharpness parameter α which we used previously.

### Model implementation

The model was implemented in MATLAB (The MathWorks Inc., Natick, MA, USA). All model computations were carried out on a Cartesian grid of 0.125 × 0.125 mm^2^ resolution. Spatial filtering used for the ODC and BOLD PSF modeling was carried out in the frequency domain using discrete Fourier transforms. The discrete Fourier transform assumes signals to be periodic, thereby forcing opposite edges of the grid to be continuous. In order to minimize this effect on modeling, the simulated area was extended relative to the data by doubling the length of each dimension.

### Prior estimation

Table 1 presents an overview of employed priors for all parameters. The spatial Gaussian white noise had an independent multivariate normal distribution with a standard deviation of 1 as its prior. Priors for the ODC model parameters were estimated from anatomical and neurophysiological data as described in the following sections.

**Table 1.** Model Parameters. Free parameters are probabilistic parameters. We used MCMC to sample from their joint distribution. Fixed parameters were estimated directly from the data and were held constant during MCMC sampling.

| Free parameters | Description | Prior | MCMC starting value |
| --- | --- | --- | --- |
| $n_{i,j}$ | Spatial white noise to be filtered, determines specific ODC pattern | Multivariate standard normal distribution | Up-sampled differential fMRI map, normalized to have a standard deviation of 1 |
| $\rho$ | Main spatial frequency of ODC | Limited normal distribution, parameters estimated from anatomical data | Set to mean of estimates from anatomical data |
| $\delta$ | Irregularity | Normal distribution, parameters estimated from anatomical data | Set to mean of estimates from anatomical data |
| $\epsilon$ | Branchiness | Normal distribution, parameters estimated from anatomical data | Set to mean of estimates from anatomical data |
| $\omega$ | Smoothness of transitions | Uniform between 0.3 and 2 based on neurophysiology data | Set to mean of limits |
| $\theta$ | Orientation of ODC | Flat | $\pi/2$ (corresponds to ODC bands parallel to medio-lateral direction) |
| $fwhm_{GE}$ | GE point-spread-function width (full width at half maximum) | Flat | 2 mm |
| $fwhm_{SE}$ | SE point-spread-function width (full width at half maximum) | Flat | 2 mm |
| Fixed parameters | Description | Estimation |  |
| $\beta_{GE}$ | GE response amplitude | Twice the median across voxels of the average between GE responses to left and right eye stimulation | |
| $\beta_{SE}$ | SE response amplitude | Twice the median across voxels of the average between SE responses to left and right eye stimulation | |
| $\hat{\sigma}_{GE}^2$ | GE measurement noise variance | Mean across voxels of estimation variance of differential GE response | |
| $\hat{\sigma}_{SE}^2$ | SE measurement noise variance | Mean across voxels of estimation variance of differential SE response | |

### Estimation of priors from cytochrome oxidase data

Four single hemisphere images of complete patterns of ODCs in the human brain taken from Adams et al. (2007) were reanalyzed. These images were originally obtained by postmortem staining for CO activity in human subjects who had lost one eye. The goal of this analysis was to find model parameters that gave rise to modeled ODC maps whose spatial power spectra most closely resemble those of real human ODCs (Fig. 2).

**Fig. 2.**
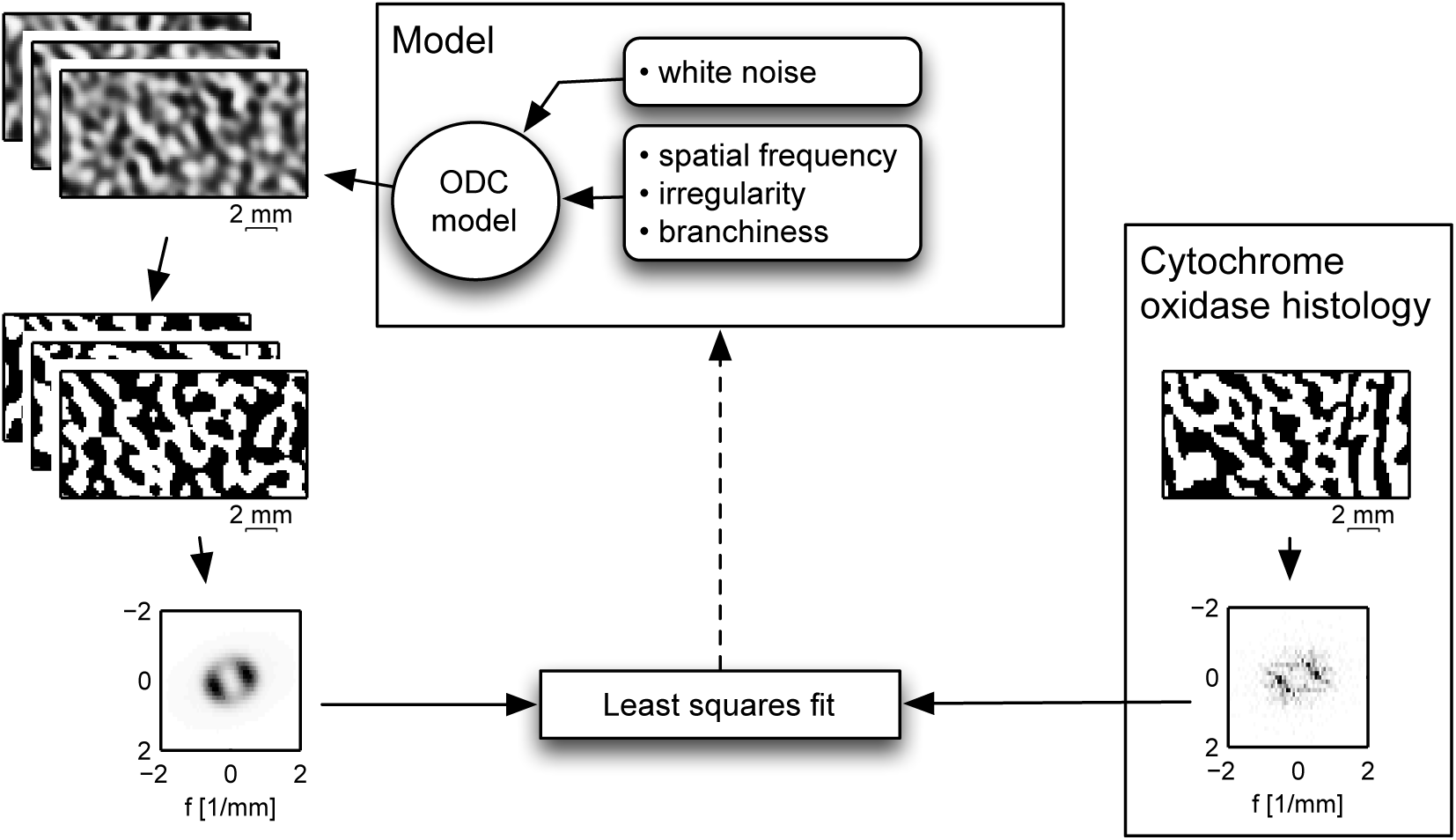
Overview of cytochrome oxidase fitting. In order to constrain the parameters of the ODC model (main spatial frequency, irregularity and branchiness), cytochrome oxidase (CO) maps of human ODCs (Adams et al., 2007) were analyzed. Model parameters were optimized so that the spatial power spectra of binary ODC maps generated by the model (left) resembled those of the CO maps (right).

Two rectangular regions from each image, which corresponded in cortical location and extent to our fMRI ODC maps, were selected. To this end, V1 boundaries and the representation of the fovea were delineated. The eccentricity for each point in V1 was computed from the cortical distance to the fovea 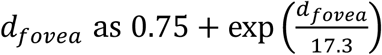 (Horton and Hoyt, 1991). For every point in the map, the two locations on the upper and lower V1 boundary (representing the vertical meridians) with eccentricity equal to that of the considered point were identified. The angular distances (along points with the same eccentricity) from the point under consideration to each of those two points on the boundary were calculated. The horizontal meridian was defined as the set of all V1 points for which those two distances were equal (green line in Fig. 3A).

**Fig. 3.**
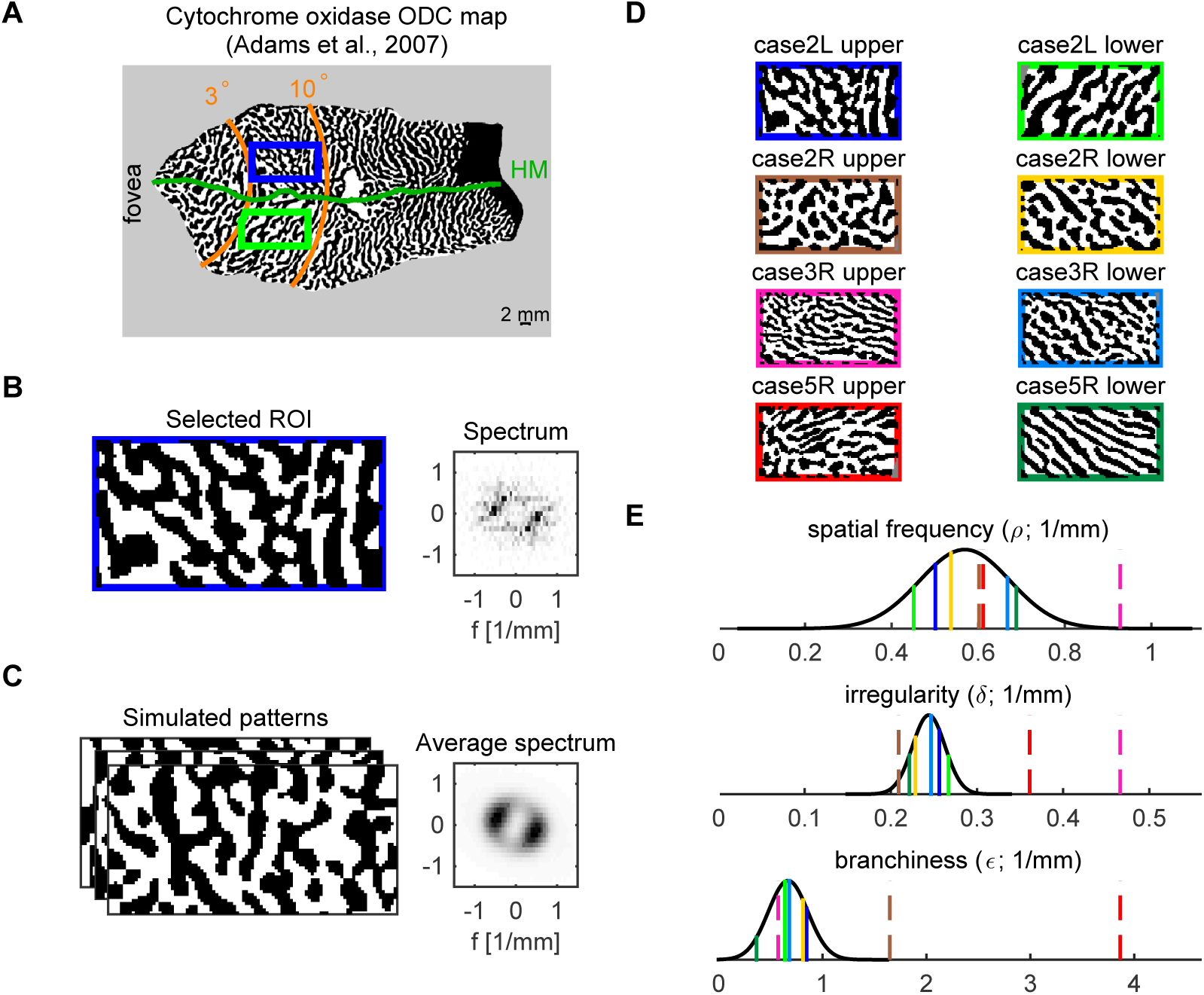
Results of cytochrome oxidase ODC map analysis. A Cytochrome oxidase ODC maps from human V1 were imported from Adams et al., (2007). Regions of interest (ROIs) were selected to be comparable in size and location to our fMRI data. B The ODC pattern from the selected upper bank region (left) is shown next to its spatial power spectrum (right). C Model parameters were optimized to produce simulated patterns (left) whose average spectrum (right) was comparable to the spectrum of the data in B. D The patterns from the upper and lower bank regions of all four cases are shown with color-coded surrounds. E The estimated values of model parameters from all patterns are shown as vertical lines using the same colors as in D. Dashed lines indicate values from patterns that were classified as outliers in the distribution of spatial frequency, irregularity, or branchiness. The mean and standard deviation of the remaining values were used to define Gaussian distributions (black) to be used as priors for fitting the model to the fMRI data.

The two regions to be selected, corresponding to the upper and lower banks of the calcarine sulcus, were then defined using the following criteria. First, the spatial extent was set to 15.7 mm x 8 mm, so that the area was equal to the mean area of our fMRI regions of interest (ROI) and the aspect ratio was equal to the mean aspect ratio of our fMRI ROIs. Second, ROIs had to be 5 mm away from the horizontal meridian and centered within an eccentricity range of 3° to 10°, corresponding to the expected location of the flat regions of the calcarine sulcus (Cheng et al., 2001).

The pattern of the CO map was binarized, in order to obtain the pattern of absolute ocular dominance (i.e. left or right eye preference). Then, for each map, we fitted the parameters of the ODC part of the model such that the spatial power spectra of the simulated binarized ODC maps were similar to those of the CO maps of ODCs (Fig. 3B, measured ODC; Fig. 3C, simulated ODC). For a model that consists of the filtering of spatial white noise only (i.e. without a sigmoidal point-wise non-linearity or binarization), the frequency spectrum of the output is expected to resemble the filter shape. Here, the spatial frequency spectra of the binarized CO maps were used to obtain first estimates of the ODC filter parameters. To that end, spectra were resampled to polar coordinates and their radial and angular averages were computed. Model equations for the radial and angular filter components (see Appendix A) were fitted to these averages using the MATLAB Curve Fitting Toolbox, enabling the extraction of parameter estimates (*ρ, δ, Θ, ∈*). Random binary ODC patterns (400 in each step) were simulated using these estimates as initial values. Their power spectra were averaged, and the sum of squared differences between the data spectrum and the average simulated spectrum was computed. An optimization algorithm in MATLAB (fminsearch; Lagarias et al., 1998) was used to find parameters that minimized this sum of squared differences.

Three maps and their fitted parameters showed outlier features. For example, ‘case5R upper’ (Fig 3D, red surround) showed very thin bands in one region (upper left) immediately adjacent to a region of thicker bands (bottom right), resulting in outlier estimates of irregularity and branchiness (red lines in Fig. 3E, middle and bottom panels). Such abrupt changes may have resulted from the processing of anatomical specimens or from the presence of curved boundaries between locally flat regions.

In order to avoid atypical parameters estimates, parameter values whose absolute deviation from the median exceeded 3.7 times the median absolute deviation (corresponding to 2.5 standard deviations of a normal distribution; Leys et al., 2013) were marked as outliers. Maps in which at least one parameter was marked as an outlier were excluded from further analysis. This procedure resulted in the exclusion of three maps (cases 2R, 3R and 5R upper) from further analysis.

For each parameter, model priors were defined as normal distributions with means and standard deviations equal to the sample means and standard deviations of parameter values from the remaining maps. In order to further discourage extreme parameter values, we set the prior for *ρ* outside two standard deviations from the mean to zero.

### Estimation of a prior for the smoothness parameter *ω*

A prior for the smoothness parameter *ω* was constructed on the basis of ocular dominance indices (ODIs) as reported in the neurophysiological literature. We assumed that ODI distributions in humans are similar to those in the macaque.

ODIs are defined as 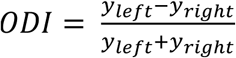, where *y_left_* and *y_right_* denote the response values to stimuli presented to the left and right eye respectively (e.g. Berens et al., 2008). ODIs were calculated from differential ODC maps generated by our model and fit to ODI distributions from the literature (Berens et al., 2008; Hubel and Wiesel, 1968). *ω* was allowed to vary while all other parameters were fixed as the mean of their anatomical data estimates. The value of *ω* that resulted in the smallest Kullback-Leibler divergence between the modeled and the target distribution was selected.

In order to fit modeled ODI distributions to the seven-class classification of Hubel and Wiesel (1968), classes 1 and 7 were collapsed into an exclusively responding class (left or right eye). Classes 2, 3, 5 and 6 were collapsed into an intermediately responding class, with class 4 responding indifferently. Modeled ODI distributions were also transformed into these three classes. The range of absolute ODIs that were assigned to the exclusively responding class and to the indifferently responding class were defined by a class width parameter. The values of *ω* and of the width parameter that together resulted in the smallest Kullback-Leibler divergence between the modeled and the target distribution were selected.

### fMRl Data acquisition

7 T BOLD fMRI data from Yacoub et al. (2007) were reanalyzed. The data were obtained from three subjects in six sessions each, using GE (three sessions) and SE (three sessions) imaging. The target ROI of one subject was unusually and densely covered by large blood vessels. We therefore excluded the data from this subject and used the two other datasets. A single slice was imaged; it was selected such that it was parallel to and maximally overlapping with a flat gray matter region of the calcarine sulcus. The in-plane resolution was 0.5 × 0.5 mm^2^ and the slice thickness was 3 mm. Each run included a baseline epoch, in which a blank gray image was presented, and alternating epochs of left or right eye stimulation. Detailed descriptions of the methods used for data acquisition can be found in Yacoub et al. (2007).

### fMRI Data processing

#### Data reconstruction

The measured k-space data were preprocessed using dynamic off-resonance in k-space (DORK) to remove respiration-induced fluctuations in resonance frequency (Pfeuffer et al., 2002). Subsequently, a Fourier transformation was applied in order to transform the data to the image space. Three datasets (subject 2, SE) were acquired using partial Fourier and were reconstructed using a homodyne reconstruction algorithm (Noll et al., 1991).

#### Motion correction

Residual head motion was corrected using AFNI’s 3dvolreg (Cox and Jesman-owicz, 1999). This algorithm requires multiple slices; therefore, identical copies of the slice were concatenated from above and below. The additional slices were later discarded from the output of the algorithm. The reference volume in each run was set to the volume with the highest average correlation to all other volumes. Each run was motion-corrected using the two-passes option and Fourier interpolation. All volumes for which any voxel was displaced more than 1 mm relative to the reference volume were marked as motion outliers and were later excluded from the general linear model (GLM) analysis. All resulting transformation matrices were saved.

Between-run motion correction was carried out by first averaging all within-run corrected volumes from each run. Next, the series created from concatenating these single-run averages was corrected in the same manner as described above. Again, all transformation matrices were saved.

Finally, the within-run corrected data was transformed by applying the saved between-run transformations. The resulting combination of two interpolations (within- and between-runs) was created for intermediate use only. For our quantitative analysis of the PSF, only one interpolation was applied. This one interpolation accounted for all alignments and registrations of the data (see below).

#### Outlier volume detection

For every volume, the measured fMRI signal in each voxel was compared to the entire time-course of that voxel by computing the z-score of the measured fMRI signal relative to the voxel’s time series across all other volumes. Volumes that were already marked as motion outliers were not included in this calculation. The volume under consideration was marked as an outlier volume if the average of z-scores across all voxels was larger than 2. Outlier volumes were later excluded from the GLM analysis.

#### Between-days and between-modality registration

ROIs for each session were imported from our previous analysis of the data (Yacoub et al., 2007). For each modality (GE or SE), between-days registration was performed using mean intensity images averaged over all runs of a given day. First, the single-day ROIs were aligned according to their centers of mass. Next, each day’s mean intensity image was cropped such that all images contained the same amount of space around their ROIs, the ROIs were in the same position, and all cropped images were of equal size. Weight masks were calculated for each day by assigning a weight of one to all voxels inside the ROI and zero to all voxels further away from the ROI than 25 mm. Voxels outside of the ROI but closer than 25 mm were assigned an intermediate weight that varied smoothly between one and zero according to the function 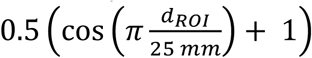, where *d_ROI_* is the shortest distance to the ROI. Out of the three days, the one whose mean intensity image had the highest average correlation to those of all other days (weighted by the mask) was selected as the reference day. Each day was registered to the reference day using FSL’s flirt 2D registration without large-scale search (Jenkinson et al., 2002; Jenkinson and Smith, 2001), using the weight masks and normalized correlation as the cost function. All transformations were saved.

Initial between-modality (GE and SE) registration was carried out using the registered GE and SE images averaged over all days. A procedure similar to that used for within-modality registration was employed, except that the correlation ratio served as the cost function. For the data of subject 1, AFNI’s 3dvolreg (using the same options as in motion correction) produced a better registration than FSL’s flirt based on visual inspection, and was therefore used

All registration results were visually inspected. Residual misalignments found in one day of subject 1 and one day of subject 2 were manually corrected.

#### Data resampling

In order to avoid smoothing of the data due to multiple interpolations, all transformation matrices (within-run motion correction, between-run motion correction, between-days within-modality registration and between-modalities registration) were combined. All unprocessed data was transformed using one single Fourier interpolation per volume using AFNI’s 3drotate (Cox and Jes-manowicz, 1999).

#### GLM analysis

For each run, a GLM was fit to each single voxel time-course. The model consisted of a constant predictor and the two stimulation paradigms (left and right eye stimulation) convolved with a standard hemodynamic response function. Volumes that were previously determined to be outliers due to extensive head motion or imaging artifacts were excluded from the fit. Relative responses were calculated by dividing the estimated stimulus response magnitudes by the estimated constant baseline. Differential responses were calculated as the difference between left and right eye responses. Unspecific responses were calculated as the average between left and right eye responses. In addition, standard errors of all estimates were calculated. For visualization purposes only, estimated response maps as well as modeled response maps were interpolated to 0.25 × 0.25 mm^2^ resolution by zero-padding in spatial frequency space and were band-pass filtered for eliminating cycles shorter than 1.25 mm and longer than 12 mm.

#### Multi-run and multi-day averaging

Standard errors were comparable between runs and days. Accordingly, single-day responses were calculated by averaging the GLM estimates of the single-run responses. Likewise, responses for each subject and imaging modality (GE or SE) were estimated by averaging single-day responses.

Standard errors for these averaged responses were estimated as standard errors of the mean from the distribution of single-day responses. Averaged response maps from all three days were used for further processing for subject 1. For subject 2, between-days correlation of a pair of SE sessions was significantly lower than those obtained from all other pairs in our data. We therefore averaged only the two most reproducible SE sessions (highest correlation of differential responses) and, separately, the two most reproducible GE sessions in order to achieve equal processing between SE and GE.

#### Optimization of between-modality registration using differential maps

Between-modality registration was further optimized. The GE and SE differential maps were shifted relative to each other vertically and horizontally by multiples of a quarter voxel up to three voxels in each direction and the set of shifts that resulted in the highest correlation between differential GE and SE maps was saved. To avoid multiple interpolations, this shift was combined with all previously found transformations (i.e. motion correction and registration) into a single transformation and interpolation. All unprocessed data were transformed over again as described above, followed by GLM analysis, multirun and multi-day averaging.

##### Quantities used for MCMC fitting

Image artifacts, noise and blood vessels may result in some voxels with extreme differential responses that would have a disproportionate effect on fitting the model. For this reason, all voxels with a differential response showing absolute deviation from the median exceeding 3.7 times the median absolute deviation (corresponding to 2.5 standard deviations of a normal distribution; Leys et al., 2013) were excluded (percentage of excluded voxels in subject 1: 5.0% GE and 4.1% SE, in subject 2: 2.8% GE and 2.3% SE). Note that although this procedure may have removed some voxels with large vessel contributions, it was not meant to systematically remove all voxels with such contributions (see discussion). We then calculated the median unspecific response and the root mean square (RMS) of the differential response standard errors from the remaining voxels. We set the maximum response amplitudes *β_GE_* and *β_SE_* to twice the median of the left/right averaged GE and SE responses, respectively, as defined by the model.

#### MCMC fitting

The posterior probability of GE and SE PSF widths given the data and priors over parameters was estimated using MCMC sampling (Fig. 1). MCMC sampling was implemented using a Hamiltonian Monte Carlo algorithm (Duane et al., 1987; see Neal, 2011 for a more recent review).

The algorithm requires input in the form of a function that computes the negative log posterior probability (the potential energy of the model; see Appendix B in the Suppl. Material) and its gradient (see Appendix C in the Suppl. Material). The log posterior probability depends on model parameter values, their prior probabilities, the data and the uncertainty of the data. The data in this sense were the maps of measured differential GE and SE responses within the ROI that were not excluded as outliers. The uncertainty of the data was characterized by the RMS of differential response standard errors, calculated separately for GE and SE. The exact form of the log posterior probability and derivations of the formulae for efficient computation of its gradient are described in the Appendix.

Two parameters determine the dynamics of parameter space exploration. The first parameter, the number of leapfrog steps per iteration, was set to a value of 20. The second parameter, the step size, was initially set to 0.005 and was adjusted adaptively so that the acceptance probability stayed close to the theoretical optimum of 0.651 (Neal, 2011). In addition, the step size was varied randomly within a range of ±20% to avoid periodicity in the trajectories (Neal, 2011).

Initial values used for all model parameters can be found in Table 1.

The MCMC algorithm was run for 512,000 iterations, of which every 256^th^ sample was retained. The set of all retained samples is an approximation to the joint posterior probability distribution of all parameters given the data and the model, while taking prior distributions into account.

See the attached video, that demonstrates the initial stages of the fitting.

##### Analysis of MCMC sampling results

The samples of PSF widths were binned into 0.04 mm wide intervals. We then identified the bin that contained the highest number of samples, which is the maximum a posteriori probability estimate obtained from the marginal distribution for the PSF width. Highest marginal posterior density credible intervals at the 95% level were computed by selecting the narrowest intervals containing 95% of the PSF width samples.

##### MCMC sampling diagnostics

The quality of the MCMC sampling process was assessed by visual inspection of parameter sample traces, autocorrelation estimates of the samples traces and the Geweke diagnostic, which is a z-test for difference between sample means in the first 10% and last 50% of samples (Geweke, 1991).

##### Estimation of T_2_/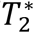 blurring or sharpening effect

Imaging modulation transfer functions (MTF) were estimated from the last volume of each run in which the phase-encode gradients were switched off (Kemper et al., 2015). This resulted in read-out lines that were expected to vary in amplitude only, according to the phase-encode direction imaging MTF (reflecting *T_2_*/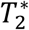 decay as experienced by all normally acquired volumes). First, the peak position along the read-out direction was found from the average of the absolute magnitudes computed over all read-out lines. Next, the imaging MTF was estimated by combining the absolute magnitudes of all read-out lines at the peak position into a vector.

We estimated the blurring or sharpening due to *T_2_*/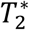 decay as a separate effect from the effect of finite and discrete MR sampling (for more details see Chaimow and Shmuel, 2016). To this end, the complex imaging PSF was computed by applying a discrete Fourier transform to the estimated imaging MTF.

Convolution of the original pattern with the real component of the complex imaging PSF is an approximation to the full MRI acquisition (Chaimow and Shmuel, 2016), including the last stage of taking the absolute of the complex values obtained at the end of the reconstruction.

We computed the inverse discrete Fourier transform of the real component of the complex imaging PSF, resulting in its MTF. Two Gaussian functions with zero means were separately fitted to the MTF of the real component of the complex PSF and to its inverse. Then, we compared the goodness of fit (R^2^) obtained by the two fitted Gaussians. If the better fit was obtained by fitting a Gaussian to the MTF of the real component of the complex PSF, the effect of *T_2_*/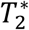 could be described as Gaussian blurring. If the better fit was obtained by fitting a Gaussian to the inverse of the MTF of the real component, the effect of *T_2_*/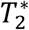 could be described as a sharpening that could reverse a specific Gaussian blurring.

We therefore computed the FWHM of the Gaussian with the better goodness of fit (obtained by fitting to either the MTF of the real component or to its inverse).

Note that in the case of sharpening, the computed FWHM characterizes the Gaussian ‘used’ for blurring which would be reversed by the sharpening effect of the *T_2_*/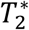 decay. FWHM estimates for each modality were first averaged over all runs of each session (day) and then over all sessions of each subject.

##### Inclusion of T2/T2* blurring in the model

A version of our model that included the effect of *T_2_*/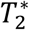 decay was fitted to our data. Depending on whether the *T_2_*/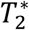 decay effect resulted in blurring or sharpening, the BOLD MTF (Appendix A, BOLD response) was changed to: 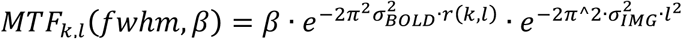 (for blurring) or 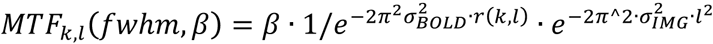 (for sharpening), where 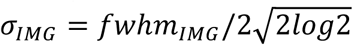 and *fwhm_IMG_* is the estimated FWHM of the Gaussian blurring kernel that models the effect of *T_2_*/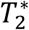 decay. We assumed the second dimension (associated with index *l*) to be the phase-encode dimension.

## Results

Our goal was to fit a probabilistic generative model to maps of ODCs obtained with GE- and SE-based BOLD fMRI (Fig. 1). We aimed to infer posterior probability distributions of model parameters, specifically the width of the GE and SE BOLD fMRI PSF.

### Parameter priors obtained from real human ODC

The ODC imaging model (Chaimow et al., 2011) consisted of simulating realistic ODCs by the filtering of spatial white noise (Rojer and Schwartz, 1990) followed by a spatial BOLD response and MRI k-space sampling.

Before we fitted our model to fMRI data, we determined priors for the ODC parameters by incorporating statistical information obtained from real human ODC patterns (Fig. 2). To this end, we analyzed CO maps of ODCs from human V1 taken from Adams et al., (2007).

It should be noted that CO labeling intensities are expected to provide a fairly accurate estimate of the preferred eye. However, there are multiple, potentially non-linear transformations between neuronal activity, staining intensity and the final processed image. These make it unlikely that the CO intensities quantitatively reflect the relative ocular dominance. Therefore, we only used binarized versions of these maps, thresholded to represent the absolute preference to either left or right eye stimulation). We eventually determined priors on ocular dominance from neurophysiological recordings (Berens et al., 2008; Hubel and Wiesel, 1968).

We first restricted the maps of the entire V1 to small regions (Fig. 3A) whose size and location were similar to those of our fMRI data (with origins in flat regions of the calcarine sulcus). Then, for each map, we fitted the parameters of the ODC part of the model such that the spatial power spectra of the simulated binarized ODC maps were similar to those of the CO maps of ODCs (Fig. 3B, measured ODC; Fig. 3C, simulated ODC). Simulated maps generated using these parameters looked qualitatively similar to the true CO maps (Fig. 3C). The set of all imported maps is shown in Figure 3D and the estimated parameters from all maps are shown color-coded in Figure 3E.

For each parameter we defined Gaussian priors that fit the distribution of all remaining parameter estimates (black curves, Fig. 3E). In particular, the prior for the main pattern frequency *ρ* had a mean of 0.57 cycles/mm with a standard deviation of 0.1 cycles/mm, which corresponds to an average column width of 0.87 mm.

### Smoothness of ODC maps

In order to construct a prior for the smoothness parameter *ω* we analyzed distributions of ocular dominance indices (ODIs) as reported in the neurophysiological literature. ODIs quantify the relative contributions of each eye to measured responses, and their distribution is tightly linked to the smoothness parameter *ω*. Small values of *ω* result in sharp transitions between columns associated with ODIs close to +1 or −1. Large values of *ω* result in smooth transitions, with few locations reaching absolute monocular responses and most ODIs being close to 0.

We analyzed ODI distributions taken from Hubel and Wiesel (1968) and Berens et al. (2008) by fitting ODI distributions computed from our model as a function of smoothness *ω*. We found a value of *ω* = 1.5 to best explain ODI distributions corresponding to the data in Berens et al. (2008), whereas the data in Hubel and Wiesel (1968) were best fitted with *ω* = 0.36. Both datasets came from macaque monkeys. Berens et al. (Berens et al., 2008) used multiunit activity, a measure whose ODIs are expected to be blurred relative to single neuron responses and are therefore expected to match a higher *ω*. Data from Hubel and Wiesel (Hubel and Wiesel, 1968) presented single-unit responses but were less quantitative. We therefore chose a uniform prior distribution for *ω*, limited by 0.3 from below and 2 from above, effectively reflecting the range of uncertainty associated with *ω*.

### GE and SE BOLD maps of ODC

Having constructed a generative model with realistic priors, the next step was to process the fMRI data and to extract all quantities needed to fit the model. We reanalyzed fMRI data from two subjects (Yacoub et al., 2007) using a general linear model (GLM) to estimate responses to left and right eye stimulation (Fig. 4). The single-eye response maps were dominated by global unspecific responses and superimposed band-shaped modulations (Fig. 4A).

**Fig. 4.**
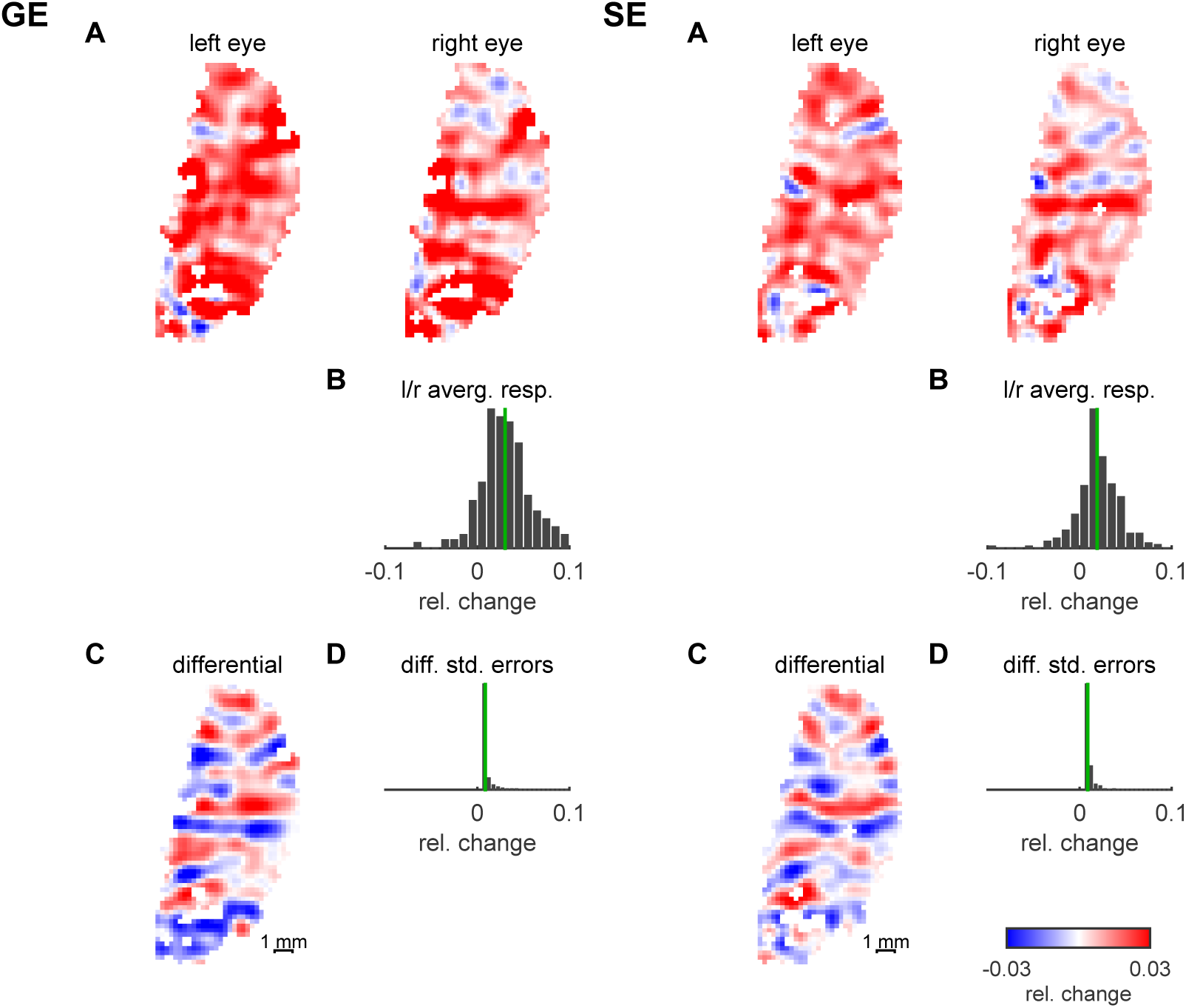
fMRI ODC data. Results from the GLM analysis of fMRI data from subject 1 for GE (left) and SE (right). **A** Responses to left and right eye stimulation relative to baseline. **B** The response maps to the left and right eyes from A were averaged. B shows the distribution of the average response. Its median (in green) was used to set the overall amplitude of the BOLD response model. **C** The difference between left and right eye responses yields the differential ODC map. **D** The distribution of standard errors of all differential responses. From this distribution we estimated the noise level used by the model. The color look-up-table applies to all response maps.

We separated these two components by first calculating the voxel-wise difference between left and right eye responses, yielding the differential ODC maps (Fig. 4C). Here, the band-shaped organization is clearly visible. The range of differential contrasts as defined by their standard deviation was 1.8% (GE) and 1.5% (SE) for subject 1, and 1.0% (GE) and 1.0% (SE) for subject 2.

In addition, we calculated voxel-wise averages of left and right eye responses (Fig. 4B). According to our model, which assumes antagonistic patterns of neuronal responses, this average response is expected to be independent of the local ocular preference. Furthermore, it is expected to be equal to a spatially homogeneous response with half the amplitude of the highest possible ocular dominance (with no response to the non-preferred eye).

We calculated the median of this left/right average response over all voxels. It was 3.0% (GE) and 1.9% (SE) for subject 1, and 3.7% (GE) and 2.0% (SE) for subject 2. In accordance with the model, we then set the amplitudes of the model PSFs to twice these values.

Finally, we estimated the measurement noise level of the differential maps as the root mean square (RMS) of all standard errors estimated by the GLM (Fig. 4D). It was 0.9% (GE) and 0.9% (SE) for subject 1, and 0.6% (GE) and 0.8% (SE) for subject 2.

#### Estimation of GE and SE point-spread widths

We went on to estimate the probability distributions of GE and SE PSF widths given our data. Theoretically, this requires integrating the posterior probability distribution of model parameters over all other parameters (including the high-dimensional spatial noise parameter). However, exact integration over this high dimensional space is not feasible. We therefore used MCMC to sample from the posterior probability distribution. Every sample contains all the parameters necessary to simulate one anatomical ODC map and the resulting GE and SE fMRI differential maps. The algorithm draws parameter samples with a probability proportional to how likely these parameters are to have generated the measured data given all the priors.

Figure 5 (second row, common ODC) shows one of many possible ODC patterns generated by our model. It was generated using the parameter sample with the highest posterior probability. Differential BOLD fMRI maps modeled as arising from this shared ODC pattern (Fig. 5 second row, model GE and model SE) resemble the data closely (Fig. 5 first row, data GE and data SE). The distribution of PSF widths from all samples (Fig. 5 bottom) is an estimate of the true probability distribution of PSF widths for that data (see the attached video that demonstrates the initial iterations of the fitting procedure).

**Fig. 5.**
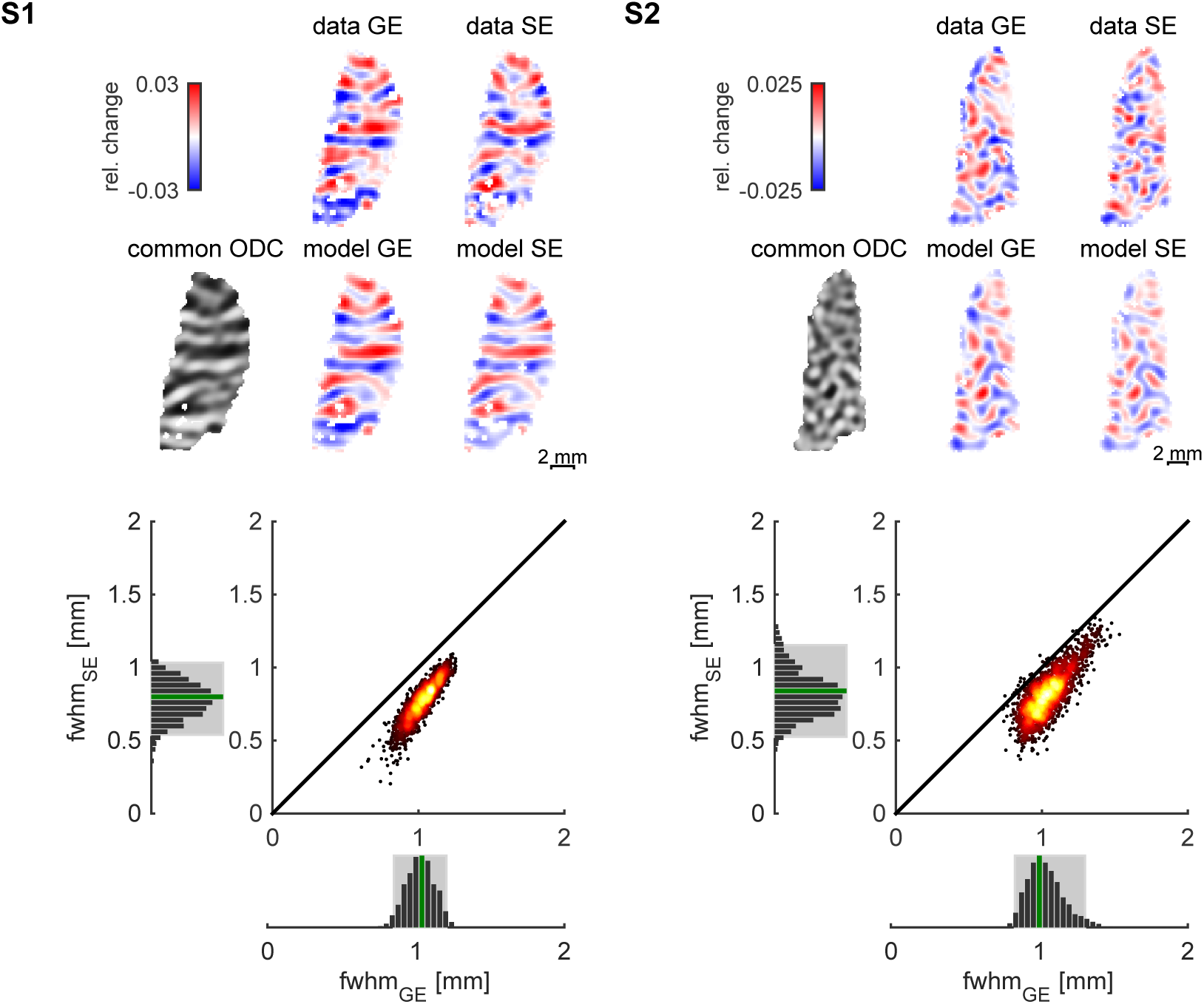
Results of point-spread width estimation. The probability distribution of PSF given the data was estimated using Markov Chain Monte Carlo Sampling. The model simulated GE and SE maps each with their own BOLD parameters and with a common underlying ODC map. Results are shown for both subjects. The first row shows the measured differential ocular dominance map from the GE (left) and SE (right) experiments. The second row shows the modeled underlying ODC maps (left) and the modeled differential fMRI maps from the maximum a posteriori (MAP) sample. The bottom part of the figure shows the joint and marginal distributions of GE and SE point-spread full-widths at half-maximum (FWHM). The gray rectangles show the 95% credible intervals (highest posterior density interval). The scatter plots show that the vast majority of individual GE PSF samples were wider than their SE PSF counterparts. The MAP estimates (green bars) obtained from the marginal distributions of the FWHMs of the PSFs were 1.04 and 1.0 mm (GE), and 0.8 and 0.84 mm (SE) for subjects 1 and 2 respectively.

**Table 2.**
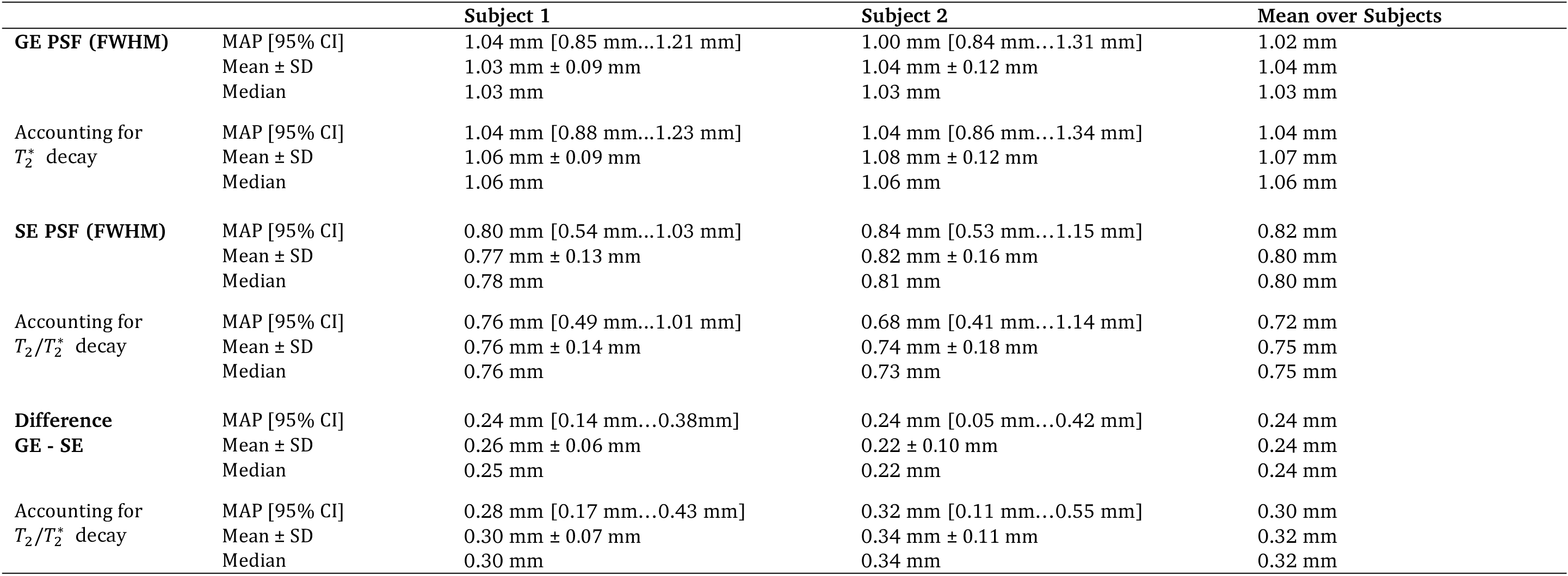
Summary of estimated PSF full-widths at half-maximum (FWHM). Maximum a posteriori estimates (MAP] with 95% credible intervals (Cl], means with standard deviations and medians were calculated from MCMC samples of the marginal posterior distributions of GE and SE PSFs FWHM and from the distribution of their differences. The results labeled with ‘Accounting for *T_2_*/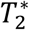 decay’ were obtained from fitting a separate model, which accounted for the effect of these decays, to the data.

Figure 5 (bottom) and Table 2 present the results of PSF widths. For subject 1, the most probable (maximum a posteriori estimate obtained from the marginal distribution of FWHMs) GE PSF width was 1.04 mm (FWHM), with 95% of the values falling between 0.85 mm and 1.21 mm. The most probable SE PSF width was 0.80 mm, with 95% of the values falling between 0.54 mm and 1.03 mm. For subject 2, the most probable GE PSF width was 1.00 mm, with 95% of the values falling between 0.84 mm and 1.31 mm. The most probable SE PSF width was 0.84 mm, with 95% of the values falling between 0.53 mm and 1.15 mm.

Furthermore, the samples of GE and SE PSF widths were correlated. This means that ODC model parameters that resulted in a relatively higher GE PSF width also resulted in a relatively higher SE PSF width. Across all modeled underlying anatomical ODC patterns, the GE PSF was almost always wider than the SE PSF. We calculated the resulting posterior distribution of differences between GE and SE PSF widths. The bottom part of Table 2 summarizes the estimated differences for the two subjects. For subject 1, the most probable difference was 0.24 mm, with 95% of the values falling between 0.14 mm and 0.38 mm. The most probable difference obtained for subject 2 was 0.24 mm, with 95% of the values falling between 0.05 mm and 0.42 mm.

### Evaluation of model fit

The validity of our results depends on how well the MCMC samples approximate the target distribution. The MCMC sampling distribution approaches the target distribution when the number of iterations goes to infinity (e.g. see Neal, 1993). For sufficiently large number of iterations, MCMC effectively samples from the target distribution.

While there cannot be proof that the target distribution has been reached, there are a number of indications that are considered reliable. The first is that the traces of samples of all parameters have settled into a stationary distribution, with no slow drifts over iterations. This can be seen in the single parameter trace plots (Fig. 6, first column) and their autocorrelation plots (Fig. 6, second column). In addition, the Geweke diagnostic (Geweke, 1991) shows that for all single parameters the mean of the first 10% of the samples was not significantly different from the last 50% of the samples (|z|<1.96). The Geweke diagnostics for the high-dimensional noise follow a standard normal distribution (Fig. 6, bottom, distribution of z-scores), as would be expected by chance under the hypothesis that the means are not different. Figure 6 also shows the dependences between PSF widths and ODC parameters (last two columns). As can be seen, higher levels of smoothness parameter (*ω*) values and to a lesser extent lower levels of the main spatial frequency parameter (*ρ*) values made a narrower PSF more likely.

**Fig. 6.**
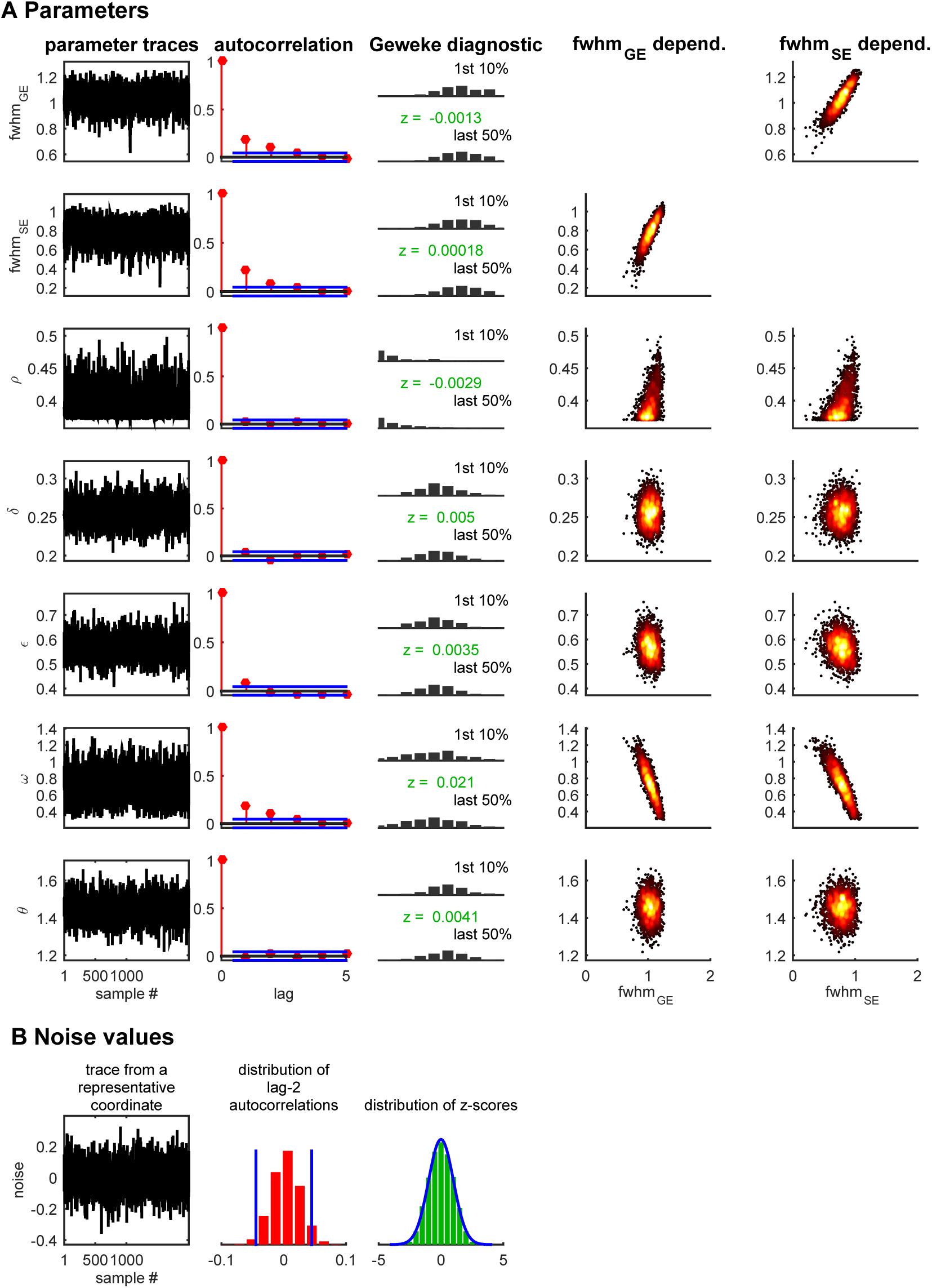
Convergence diagnostics of Markov Chain Monte Carlo sampling. Markov Chain Monte Carlo needs to run for a sufficient number of iterations in order to yield samples from the modeled probability distribution. Indications for convergence are: (1) stationarity of the parameter sampling distributions, and (2) sample autocorrelations decrease rapidly with increasing lag, relative to the total number of samples. This figure examines convergence for subject 1. The upper part of the figure (A) shows diagnostics for the standard model parameters. The bottom part (B) shows diagnostics for the white noise values that act as parameters to determine the ODC pattern. The first column (A and B) shows traces of the sampled parameters. For the noise values (B), one exemplary trace is shown from the center of the map. The second column (A and B) shows sample autocorrelations as a function of lag. The horizontal blue lines (A) indicate the 95%-confidence bounds around 0 for a white noise process. Consecutive samples (lag=1) show low autocorrelation. However, samples of lag 2 (and higher) show autocorrelation estimates that are comparable to those obtained from uncorrelated white noise. In B, the autocorrelation from the noise samples are summarized by the histogram of lag-2 autocorrelations from all coordinates together. Here, 95%-confidence bounds for a white noise process are indicated by vertical blue lines. The third column (A and B) presents the Geweke convergence (stationarity) diagnostic, which is a z-test (z-scores shown in green) for testing whether the means of the first 10% and last 50% of samples are different. In A, 2 histograms per each parameter show how similar their respective distributions are. In B, the z-scores from the noise samples are shown as a histogram together with a blue plot of the standard normal probability density representing the null-hypothesis of z=0. The last two columns (A) show the sample covariation between each parameter (vertical axis) and the GE and SE point-spread function FWHM (horizontal axis).

## Discussion

### Possible confounds: our estimates are upper bounds of BOLD fMRI spatial specificity

The PSF widths that we estimated (1.02 mm for GE BOLD, 0.82 mm for SE BOLD) reflect the realistically achievable spatial specificity of BOLD signals at ultra-high field strength (7T). However, they are only upper bounds for the true BOLD PSF widths. Subjects’ head motion, data interpolation and intra-acquisition *T_2_*/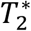 decay can all introduce additional blurring (but see section below on the effect of *T_2_*/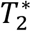 decay), causing the estimated PSF to be wider than the true PSF.

In order to minimize head motion, data was acquired from trained subjects using a bite bar. Before each scan, the position of the region of interest (ROI) was checked and the slices repositioned if necessary. We corrected the data for residual head motion and discarded any problematic volumes. We aligned data from multiple days and checked the alignment carefully. In order to further optimize between-modality registrations, we also took the differential fMRI response patterns into account, making use of the fact that they emerged from the same underlying neuronal ODC pattern.

Motion correction and between-day registration required spatial interpolation of the data. We minimized any blurring effects by applying all spatial transformations combined using one single Fourier interpolation (Cox and Jesmanowicz, 1999).

All high spatial resolution BOLD fMRI experiments will be influenced by these effects to a similar degree as ours, making our reported PSF widths good estimates for the practically relevant compound effect.

### Possible confounds: the contribution of the imaging PSF to the total BOLD fMRI PSF

In addition to the effects of the hemodynamic and metabolic responses on the spatial specificity of fMRI, the MRI acquisition process influences the effective resolution of the acquired images. Specifically, the sampling of k-space by means of temporal gradient encoding defines the spatial resolution. However, the effective spatial resolution along the phase encoding direction in EPI acquisitions can be subject to blurring or sharpening, because of *T_2_*/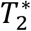 decay while the k-space is being sampled. This can potentially contribute to the overall measured spread of the BOLD fMRI signal.

In order to minimize this effect, our data were acquired using a reduced field-of-view (in SE) and multiple segments. These measures limited the total read-out duration per segment (25.6 ms for GE and 24 ms for SE) to approximately the 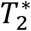 of the tissue (Uludağ et al., 2009) and are expected to result in only minor blurring or sharpening (Haacke et al., 1999)).

We estimated the blurring or sharpening and their contributions to the total BOLD PSF. In general, MRI data acquisition using EPI has two distinct effects on the effective spatial resolution. The first is the effect of the finite and discrete MR sampling with no decay. However, in the current study MR sampling was part of the model and therefore has already been accounted for. The second effect is the already mentioned *T_2_*/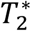 decay blurring or sharpening. This effect is limited to the phase encode direction (vertical direction in all presented maps). In order to characterize it, we estimated the imaging modulation transfer functions (MTF) along the phase encoding direction from a reference volume obtained in each run, in which the phase-encode gradients were switched off (Kemper et al., 2015).

Using a model of Gaussian convolution and MRI sampling (Chaimow and Shmuel, 2016, in preparation) we obtained Gaussian functions that can model the separate effect of *T_2_*/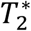 decay. We then used MCMC to fit a separate version of our model to our data, where we applied the decay effect to the simulated ODC patterns by modulating the values acquired in the simulated k-space. The results presented in Table 2 show that the effect of the signal decay on the total BOLD PSF was small. For GE, while accounting for the signal decay, we obtained PSFs wider than the effective PSF obtained directly from the BOLD fMRI response. This indicates that signal decay in the GE fMRI used for obtaining our data has a sharpening effect. In contrast, the signal decay in the SE fMRI used for obtaining our data has a blurring effect. These results demonstrate that the physiological BOLD response measured with GE fMRI (that theoretically does not include signal decay) is less spatially specific than the same physiological BOLD response measured with SE fMRI (with no signal decay). This difference in spatial specificity of the GE and SE BOLD responses (with no signal decay) is even slightly larger than the corresponding effective difference obtained from the overall measured fMRI responses with signal decay.

### What do our estimated point-spread function widths describe?

The BOLD PSF describes the spatial specificity of the BOLD fMRI signal by characterizing the spatial response that would be elicited by a small point stimulus. Specifically, our BOLD PSF width measures the spread of the BOLD fMRI response (I) elicited by a small spot of neuronal activity, (II) along the cortical manifold, (III) using a differential response analysis, (IV) assuming that in a differential analysis paradigm the average spread can be described by a Gaussian function, and (V) considering a relatively long time scale.

#### (I) BOLD PSF relative to the local neuronal activity

To the best of our knowledge, our PSF estimates are the first to quantify the BOLD spread in human subjects relative to local neuronal activity. We previously estimated the FWHM of the 7 T GE BOLD PSF to be smaller than 2 mm by measuring the spread of the BOLD fMRI response around the V1 representation of edges of visual stimuli (Shmuel et al., 2007). We expect that our previous estimates as well as others’ (Engel et al., 1997; Parkes et al., 2005) included contributions from non-zero extent of receptive fields and the scatter of receptive field position of neurons in V1.

Hubel and Wiesel (1974) reported that in the macaque “… a 2 mm × 2 mm block of cortex contains the machinery needed to analyze a region of visual field roughly equal to the local field size plus scatter””.

These observations suggest that visual stimuli will result in neuronal activity that is blurred on the surface of human V1. All PSF widths that have been estimated using spatial representations of visual stimuli included this neuronal spread by nature of their experimental design. In the current study, we instead used a spatial structure of neuronal responses that is inherent to the cortex— ODC patterns. This allowed us to estimate a PSF that does not contain contributions from the spatial spread of responses to visual stimuli.

There are a number of measures of neuronal activity that a BOLD PSF could potentially relate to, notably single-unit activity (SUA), multi-unit activity (MUA) and local field potentials (LFP). Under specific circumstances, these measures can show very different activity. Under most conditions, however, they are highly correlated. This is likely to be true when mapping a cortical columnar organization. The main difference is that the spatial extent (that influences the smoothness of the spatial response pattern) of these signals increases from SUA to MUA to LFP. We estimated a smoothness prior using ODI distributions of SUA and MUA. Consequently, our PSF is based on these signals. The BOLD PSF from LFP would be narrower than our estimate because of the wider cortical spread of LFP compared to MUA activity (Xing et al., 2009).

#### (II) Spatial BOLD response along the cortical manifold

It has been demonstrated (Polimeni et al., 2010) that the PSF consists of different radial and tangential components relative to the cortical surface. The radial component describes the spread across cortical layers while the tangential component describes the spread parallel to the cortical surface. Here we investigated the tangential PSF, averaged over all layers. This is the component that is most relevant for imaging the representation of cortical columns parallel to the cortical surface. Accordingly, the location and orientations of voxels, the ROI, and the voxel size were all optimized to sample gray matter tangentially and to obtain an average from all layers.

It should be noted that there are some differences in cerebrovascular organization with respect to radial and angular direction (Duvernoy et al., 1981). The largest blood vessels are the pial surface veins that extend in various orientations along the tangential plane. Somewhat smaller are cortical-penetrating veins that are organized radially, traversing the different cortical layers. The smallest vessels, the capillaries, form a fine mesh that locally appears to be isotropic. However, their density varies with cortical layers (Weber et al., 2008). For these reasons, we cannot directly apply our PSF to the imaging of cortical layers. In addition, the distinctiveness and finite extent of layers appear to make a PSF convolution model ill-suited for fMRI of cortical layers. However, some recent results (De Martino et al., 2015; Fracasso et al., 2016; Muckli et al., 2015; Olman et al., 2012) suggest it is possible to differentially resolve layer-specific signals on the scale of 1 mm or less.

#### (III-IV) On modeling the average differential BOLD response as a Gaussian PSF

We assumed the average (over space) PSF to be a Gaussian function. However, the shape of the spread in specific cortical locations may be more complex and location-dependent (Kriegeskorte et al., 2010; Polimeni et al., 2010). Also, its width as well as the magnitude of the response may vary due to local variations in vascular geometry. In fact, the relatively wide distribution of average responses in our data (Fig. 4, distribution of l/r averg. resp.) supports this latter intuition. Therefore, a convolutional model with a single Gaussian function can only be an approximating simplification. Nevertheless, we believe that such a simplifying approach provides a useful approximation for planning and interpretation of high-resolution fMRI studies and for quantitative modeling.

As part of our pre-processing before fitting the model to the data, we removed 1.5 - 5.0 % of voxels that had extreme differential values (see methods section for precise criterion for exclusion). Part of these voxels were located in areas that were previously shown to contain blood vessels (Shmuel et al., 2010). However, for our current analysis we did not explicitly and systematically remove voxels that were affected by larger blood vessels. Our reasoning was that a consistent removal of all voxels suspected to be influenced by larger blood vessels would have reduced contiguous areas of ODCs, which would have made the model fitting more difficult.

We expect the influence of geometric variations in local vasculature to be higher for veins and venules than for capillaries because of their respective diameters and densities. Consequently, the GE BOLD signal, which is more sensitive to larger pial surface veins will be more affected by these local variations. As a result, GE BOLD imaging does not only suffer from a slightly wider PSF than SE BOLD, but it is also subject to local distortions when larger blood vessels are present.

However, although draining veins may show responses with a preference to a subset of features encoded in a columnar organization (Shmuel et al., 2010), differential analysis reduces contributions from macroscopic vessels because of their tendency to drain blood from a region larger than that of a small number of columns. Taken together, a Gaussian PSF model by itself is likely not a good model for single-condition imaging when influenced by large blood vessels (e.g. in GE BOLD imaging). In contrast, we expect that a Gaussian PSF is a good model in a differential analysis paradigm, which reduces contributions from macroscopic vessels. The BOLD PSFs we report here reflect the spatial specificity that can be achieved in a *differential* paradigm. They do not reflect the spatial specificity expected from *single-condition* imaging that involves contributions from macroscopic vessels, such as single-condition GE fMRI and to a lesser extent, single-condition SE fMRI.

#### (V) Spatial specificity as a function of stimulus duration

It has been shown that the early phase of the positive BOLD response (up until ∼4 s after stimulus onset) is spatially more specific than the later phase (Goodyear and Menon, 2001; Shmuel et al., 2007). On the other hand, stimulation paradigms that use very brief stimulation durations suffer from a highly reduced contrast-to-noise ratio, because the response does not develop to its highest potential amplitude.

We found previously that after 4 s the spatial BOLD response remained stable and that the entire spatiotemporal response could be well approximated by the first spatial principle component (Shmuel et al., 2007). Aquino et al. (2012) modeled the BOLD response as a travelling wave evolving in time and found that deconvolution of neural dynamics using such a model resulted in physiologically more plausible spatiotemporal patterns than when using a model separable in space and time (Aquino et al., 2014). The spatial profile alone, however, was very similar for both models.

Taken together, long stimulation paradigms are an efficient way of highresolution imaging and their spatial PSF can be well described by a single time-independent component. The stimulation periods for our data were 48 s long, thereby making our PSF most applicable to long stimulation paradigms.

### Spatial specificity of the BOLD response

#### Constraints on the spatial specificity of BOLD

The positive BOLD signal depends on decreases in deoxyhemoglobin content in the capillaries which then propagate downstream to draining venules and veins. These decreases are caused by elevated cerebral blood flow (CBF) and only smaller fractional increases in the oxygen consumption rate, following increases in neuronal activity. CBF is regulated at a sub-millimeter scale: (Duong et al., 2001). Similarly, Vazquez et al. (2014) reported a spread of cerebral blood volume (CBV) of 103 - 175 μm (FWHM) in mice using optical imaging. Although this measure is not directly comparable to the CBF spread in a different species (human subjects), it demonstrates that hemodynamic signals can show very high spatial specificity. The CBF response is the ultimate lower limit for the spatial specificity of any BOLD-based technique.

The deoxyhemoglobin content changes in the draining venules and veins are ultimately diluted downstream, because the draining veins pool blood not only from active but also from non-active regions. For an activated area of 100 mm^2^, Turner et al. (2002) estimated the maximal extent of undiluted oxygenation changes along a vein to be 4.2 mm. For these reasons, we can expect the PSF width of any BOLD-based imaging technique to fall in this range; that is, less than 1 mm (Duong et al., 2001) to approximately 4.2 mm (Turner, 2002). The values will be determined by how much weighting towards the microvasculature can be achieved and on the actual presence of larger draining veins in the region of interest.

#### PSF dependence on field strength

At standard magnetic fields, the width of the BOLD PSF has been estimated to be 3.5 mm for 1.5 T GE BOLD (Engel et al., 1997), 3.9 mm for 3 T GE BOLD and 3.4 mm for 3 T SE BOLD (Parkes et al., 2005). These estimates of PSF widths were confounded by the above described receptive field and scatter effects. We can make a rough estimate of what the non-confounded PSF widths at lower fields would be. We assume that on average the receptive field effect can be modeled as another convolution with a Gaussian. It follows that the square of the confounded PSF width is equal to the sum of squares of the receptive field effect width and the non-confounded PSF width. For the receptive field effect we get an FWHM of 2.12 mm when using 2.35 mm as the 7 T GE BOLD confounded PSF width (Shmuel et al., 2007) and 1.02 mm as the corresponding non-confounded PSF width (results from our current study). This in turn results in non-confounded estimates of 2.8 mm (1.5 T GE BOLD), 3.3 mm (3 T GE BOLD) and 2.7 mm (3 T SE BOLD).

These PSF widths are considerably larger than the estimates from the current study (1.02 mm for 7T GE BOLD, 0.82 mm for 7T SE BOLD). The reason for this is that the BOLD signal (both GE and SE BOLD) at lower field strengths is dominated by intravascular signals from draining veins (Jochimsen et al., 2004; Uludağ et al., 2009). At higher field strengths, the contributions from intravascular signals are reduced due to a shortening of the venous blood *T*_2_. In parallel, the relative contributions of extravascular signals around small vessels increase (Duong et al., 2003; Uludağ et al., 2009; Yacoub et al., 2003; 2001).

All PSF widths from field strengths of up to 3 T appear to fall close to the wider end of possible PSF widths. In contrast, PSF widths using SE and GE at 7 T appear close to their theoretical minimum.

#### 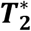 and **T**_2_ based imaging methods: GE, SE and GRASE

We found the SE BOLD PSF to be narrower than the GE BOLD PSF. This is expected because the refocusing pulse in SE imaging suppresses the extravascular signal around larger blood vessels while leaving the signal around the microvasculature intact. As a result, compared to GE BOLD fMRI, SE BOLD signals obtained at 7T have relatively larger contributions from the spatially more specific microvasculature, whereas at lower field strength the signal of either SE or GE BOLD fMRI is dominated by intravascular contributions of large blood vessels.

However, the suppression of extravascular signal around larger blood vessels by SE at high fields is not perfect. Only the k-space data that is sampled at the exact echo time will result in absolute suppression (pure *T_2_* weighting as compared to 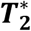 weighting). The extent to which sampled k-space data is affected by 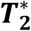 weighting increases with increasing total read-out time. Consequently longer total read-out times in SE result in decreased spatial specificity (Goense and Logothetis, 2006) and are expected to have a wider point-spread function (though still narrower than GE).

Other *T_2_* based functional imaging methods such as GRASE (Oshio and Feinberg, 1991) and 3D-GRASE (Feinberg et al., 2008) are expected to have similar spatial specificity as SE. Whether their PSFs are slightly wider or narrower will mainly depend on the 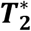 weighting component associated with such methods (i.e. echo train lengths of gradient recalled echoes employed in be-tween successive 180° pulses), in addition to their *T_2_* component. In fact, Kemper et al. (2015) have reported that 3D-GRASE had a smaller bias towards pial surface veins owing to the smaller 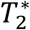 contribution when a reduced field of view is employed in zoomed 3D-GRASE compared to the longer in-plane echo-train of 2D-SE EPI.

Although we found a wider PSF for GE BOLD than for SE BOLD fMRI, the difference was relatively small (1.02 mm for GE BOLD, 0.82 mm for SE BOLD). We believe that this is due to the fact that the influence of larger blood vessels can be reduced by using a differential imaging paradigm, even when using 7T GE BOLD fMRI. Consequently, both GE and SE BOLD imaging techniques seem capable of resolving cortical columns when applying differential imaging analysis.

However, GE maps are more susceptible to confounds introduced in voxels containing blood vessels which may not be fully suppressed in differential imaging. Therefore, obtaining results of high spatial specificity using GE depends on the region of interest and on methods to mask out blood vessels.

SE is less susceptible to large-vessel confounds, that may not be suppressed by differential imaging. The response amplitude of SE is lower than that of GE. However, for imaging of highly granular structures such as ODC’s at such high resolutions, the *differential* contrast is similar for GE and SE fMRI. Overall, we believe that SE is the method of choice for mapping finer structures, especially when relying on single-condition analysis. However, which data acquisition method is optimal depends on the goal of the study and the spatial scale of the neuronal architecture under investigation.

### The application of probabilistic models of cortical columns and MR imaging

We have extended our quantitative model for imaging ODCs to a probabilistic generative model and used it to infer the PSF widths by means of MCMC sampling.

A critical component to the successful application of MCMC to our model is the Hamiltonian Monte Carlo (HMC) algorithm (Duane et al., 1987), which makes use of the gradient of the model posterior probability. Importantly, we were able to derive an efficient way to compute this gradient (Appendices C and D). HMC has the advantage of very efficiently exploring the parameter space. However, for high-dimensional problems such as ours, every step may take a long time because the gradient components for all variables need to be computed. Because of the specific form of the computations in our model (convolutions and a point-wise non-linearity), it was possible to compute the gradient efficiently as a combination of convolutions and point-wise nonlinearities as well. In principle, such efficient computation should be possible for a wide range of similar models, making HMC a powerful method for fitting such models.

We believe that the novel approach we introduce to the field of imaging cortical columns, of fitting a model of imaging columns to corresponding measured data, will be useful beyond our current study. For example, when imaging an unknown columnar structure, questions about its organization (e.g. isotropy, spatial frequency, irregularity) can be addressed via inference on model parameters.

## Conclusion

We have quantified the BOLD PSF in human subjects relative to neuronal activity, avoiding the confounding effects of scatter and size of visual receptive fields which were not eliminated in previous estimations (Engel et al., 1997; Parkes et al., 2005; Shmuel et al., 2007). As a result, our BOLD PSF estimates characterize the spatial specificity when employing imaging of fine scale cortical organizations such as cortical columns. Previous studies have shown that BOLD fMRI at 4 T and 7 T is capable of resolving cortical columns on the submillimeter scale when differential analysis is employed (Cheng et al., 2001; Menon and Goodyear, 1999; Yacoub et al., 2008; 2007; Zimmermann et al.,2011). Our results provide a quantitative basis for this resolvability and facilitate planning and interpretation of high-resolution fMRI studies of fine scale cortical organizations.

## Acknowledgements

This work was supported by grants from the Natural Sciences and Engineering Research Council of Canada (AS, NSERC Discovery grants RGPIN 375457-09 and RGPIN-2015-05103).

## Appendix A. Model of imaging ocular dominance columns

### Preliminaries

Simulations before MRI sampling were carried out on a grid of size 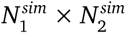. Simulations of MRI data were of size 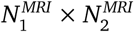. We use *i*, *j* and *k, l* as indices of 2-dimensional image and spatial frequency space, respectively. Furthermore, *r*(*k*, *l*) is the absolute spatial frequency and *ø*(*k, l*) the orientation that the point with indices *k*, *l* represents. The two-dimensional discrete Fourier transform and its inverse (D.3 and D.4) are denoted as dft2 and idft2.

### Overview over model computations

The ocular dominance columns (ODCs) imaging model can be described as a function

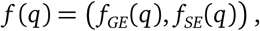

that takes the the list of model parameter values *q* = ((*n*_i, j_), *ρ*, *δ*, *∈*, *θ*, *ω*, *fwhm_GE_, fwhm_SE_*) and the fixed parameters *ß_GE_* and *ß_SE_* (see Table 1) as input and generates differential fMRI maps of ODC 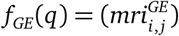 and 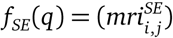 in a number of steps:

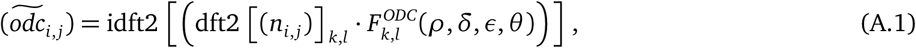

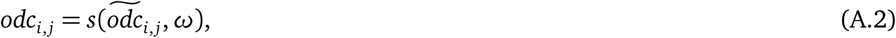

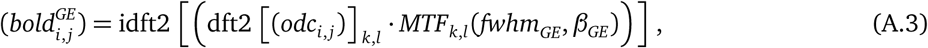

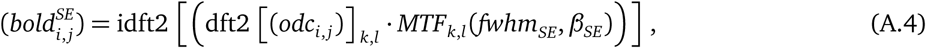

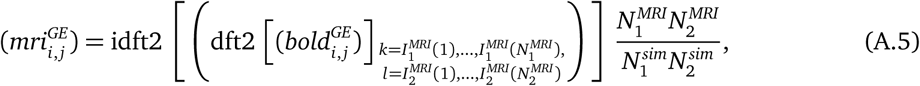

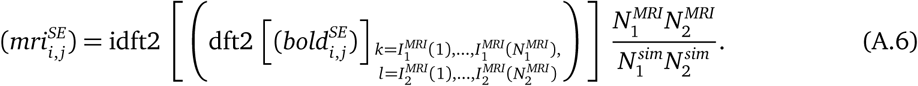

(A.1): The ODC pattern (*odc_i,j_*) was modeled by filtering two-dimensional Gaussian white noise (*n_i, j_*) using a non-isotropic filter 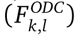 and followed by (A.2) a point-wise sigmoidal non-linearity s that controlled the smoothness of transitions between left and right eye preferrence columns. (A.3 and A.4): The BOLD response was modeled as a convolution with a Gaussian point-spread function. It was implemented as multiplication in spatial frequency space with its Fourier transform the modulation transfer function (*MTF_k>l_*). (A.5 and A.6): MRI sampling was simulated by restricting the spatial frequency space representation to its central part (indices given by index functions 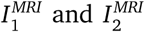) in accordance with the voxel size. The last factor corrects for the reduction in scale caused by applying idft2 to the reduced grid size.

We can combine operations A.3 and A.5, as well as A.4 and A.6:

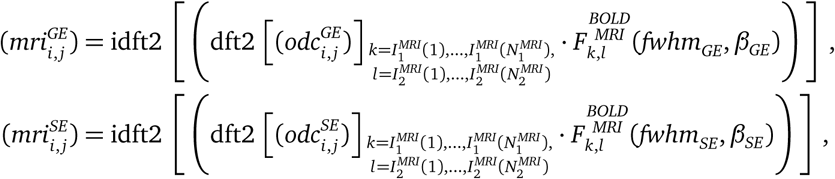

where 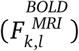is of size 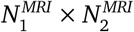 with:

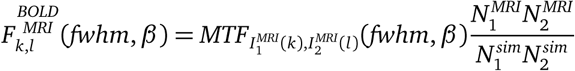

### ODC filter

An unnormalized ODC filter 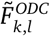 was defined in spatial frequency space as the product of radial and angular components:

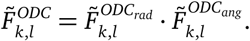

The radial component 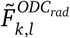 is the sum of two Gaussian functions centered on *+ρ* and —*ρ*, where *ρ* is the main spatial frequency of the pattern.

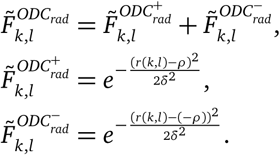

The angular component is the sum of two von Mises distribution functions centered on +*θ* and — *θ*, where *θ* is the orientation of the pattern:

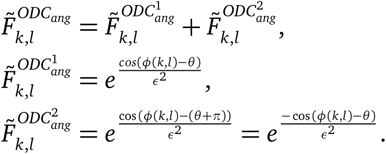

In order for the filter output to have the same variance as the input (independent of filter parameter values) we normalized the filter:

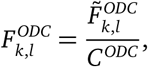

where

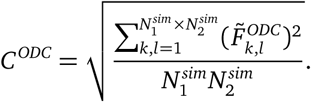

### Sigmoidal non-linearity

The point-wise sigmoidal non-linearity *s(x*, **ω*)* was defined as:

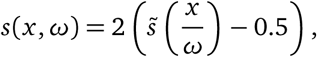

with the standard sigmoidal function defined as:

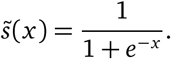

### BOLD response

The BOLD response modulation transfer function was defined as

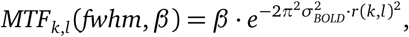

with:

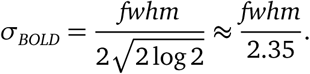

It is the Fourier transform of a Gaussian point-spread function with a full-width at half-maximum of *fwhm*. It is scaled such that a spatially extended neuronal response of 1 results in a BOLD response of amplitude *β*.

### MRI sampling

MRI sampling was simulated by restricting the spatial frequency representation according to the following index functions:

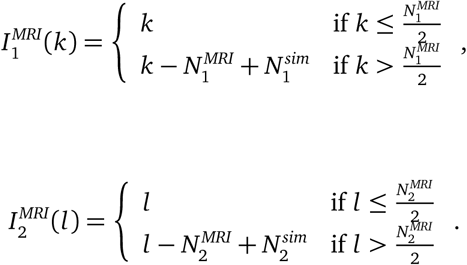

## Appendix B. Posterior probability and potential energy

### Posterior probability

Let 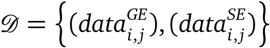 be the data of differential fMRI maps imaged using GE and SE BOLD fMRI, respectively. The likelihood - the probability to observe the data *D* given a specific set of model parameter values *q* is:

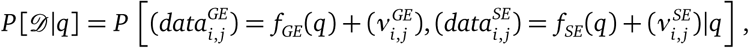

where 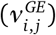 and 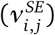 are patterns of measurement noise.

We assume the measurement noise to be independent between voxels and imaging modalities and to be distributed normally with (estimated) variances 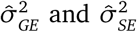. Furthermore we define 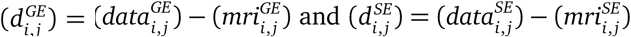 to be the patterns of deviations of the data from the model prediction. The likelihood can then be expressed as:

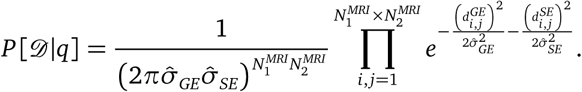

From this result we can calculate the posterior probability of parameters *q* given the data *D*:

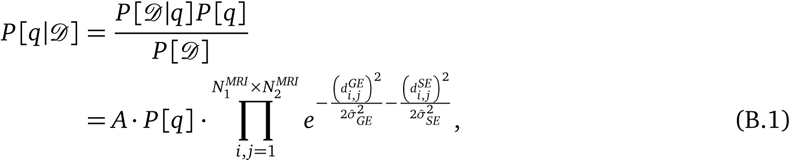

where 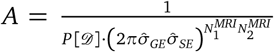 is a constant factor that is independent of *q* and *P*[*q*] is the prior probability over parameters.

### Prior probability

The prior probability over all parameters *q* was defined as:

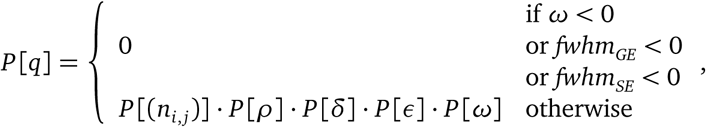

with individual parameter priors were defined as:

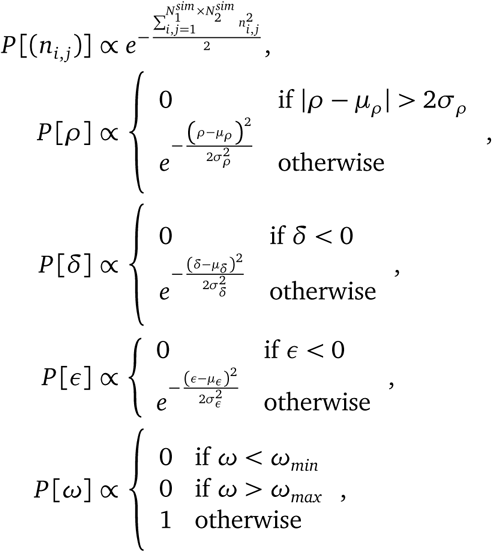

where *μ_ρ_*, *μ_δ_*, *μ_∈_* and *σ_ρ_*, *σ_δ_*, *σ_∈_* are the means and standard deviations of the ODC priors that were estimated from cytochrome oxidase data and *ω*_*min*_ and *ω*_*max*_ are lower and upper limits for **ω** that were set based on results from the neurophysiological literature.

### Potential energy

The potential energy of a state of parameter values *q* is its negative log-posterior probability plus an arbitrary constant *C* :

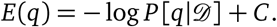

We apply B.1 and express the energy as the sum of three parts:

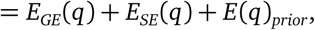

with:

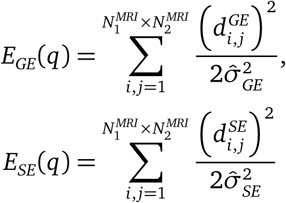

and

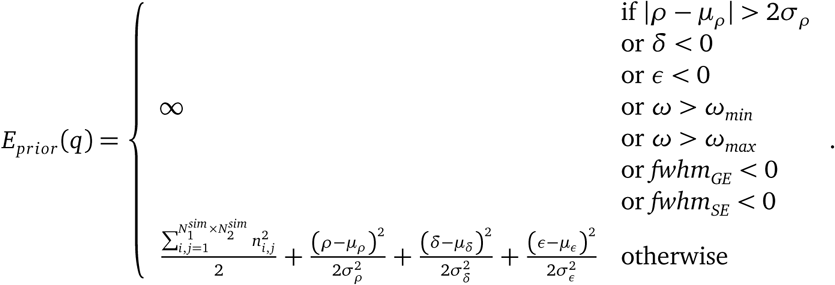

## Appendix C. Potential energy gradient

We start by deriving derivatives for functions used by the model.

### Derivatives of the ODC filter

The derivatives of the unnormalized radial and angular filter component parts with respect to their prarameters are:

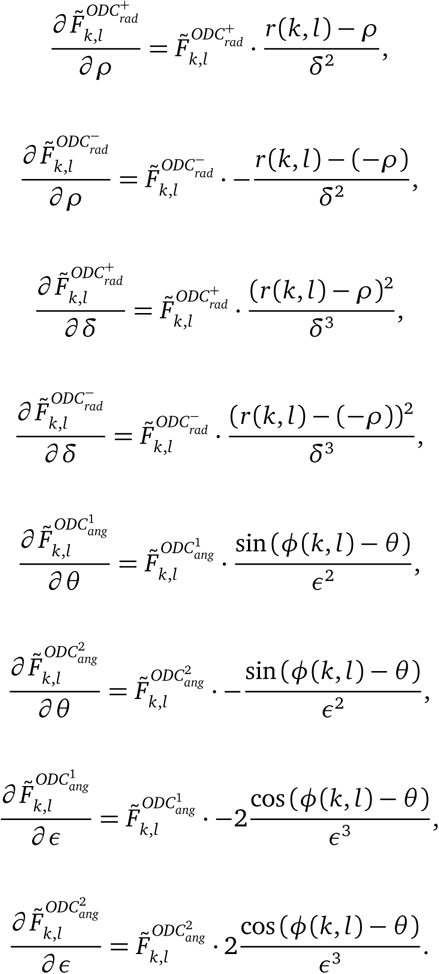

We combine these derivatives to form derivatives of the full radial and angular components:

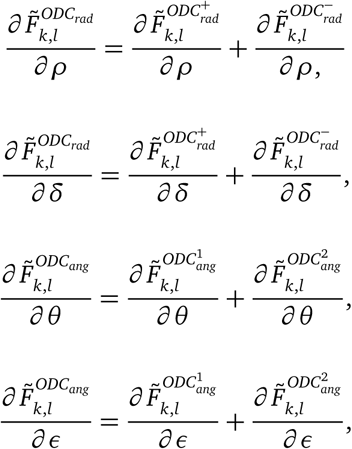

and in turn to form the derivatives of the full unnormalized filter:

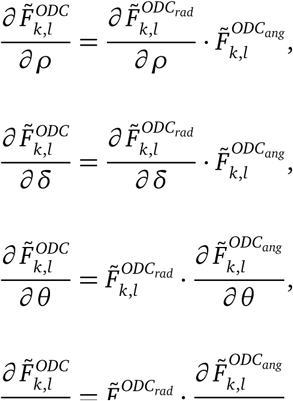

The derivatives of the normalization constant are:

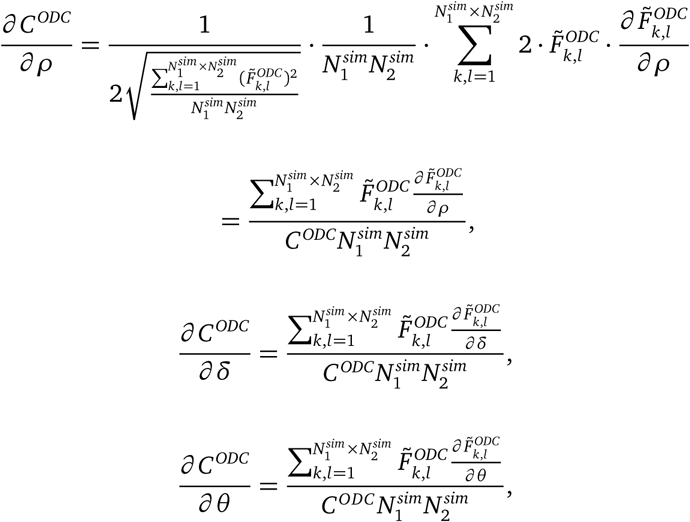

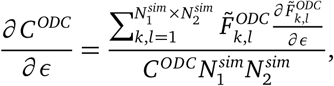

resulting in the following derivatives of the complete normalized filter:

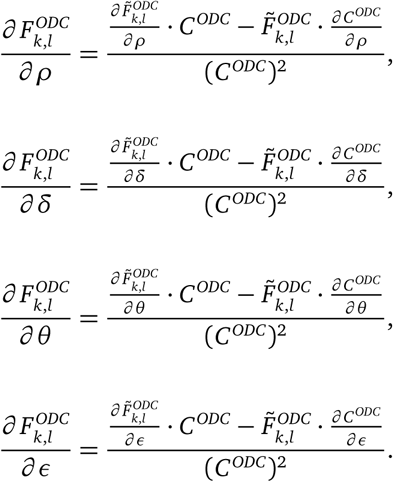

### Derivatives of the sigmoidal non-linearity

The derivative of the standard sigmoidal function 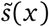 is:

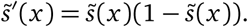

Using this result we get the following derivatives for our sigmoidal non-linearity *s*(*x*):

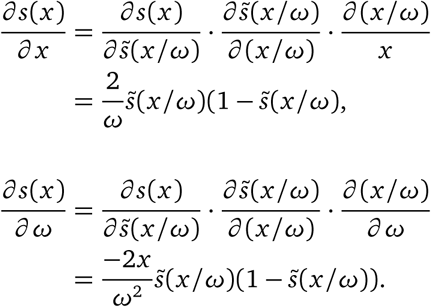

### Derivative of the BOLD modulation transfer function

The derivative of the BOLD modulation transfer function is:

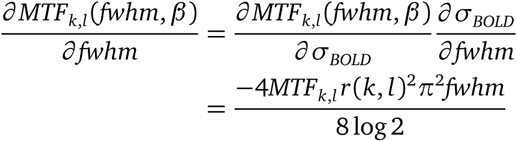

The derivative of the combined BOLD-MRI filter is:

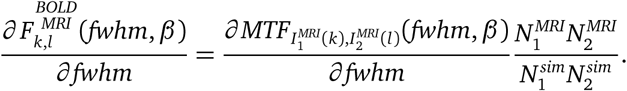

### Potential energy gradient components

The energy gradient is composed of the following derivatives:

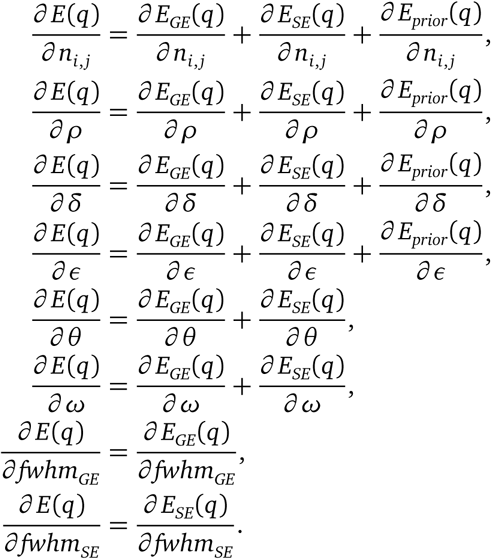

The gradient is not defind for |*ρ* - *μ_*ρ*_*| > 2σ*_*ρ*_* or *δ* < 0 or *∈* < 0 or *ω* < *ω*_*min*_ or *ω* > *ω*_*max*_ or *fwhm_GE_* < 0 or *fwhm_SE_* < 0 (regions were *E_prior_*(*q*) = ∞).

### Energy gradient with respect to noise variables

The derivatives of the *GE* energy component are:

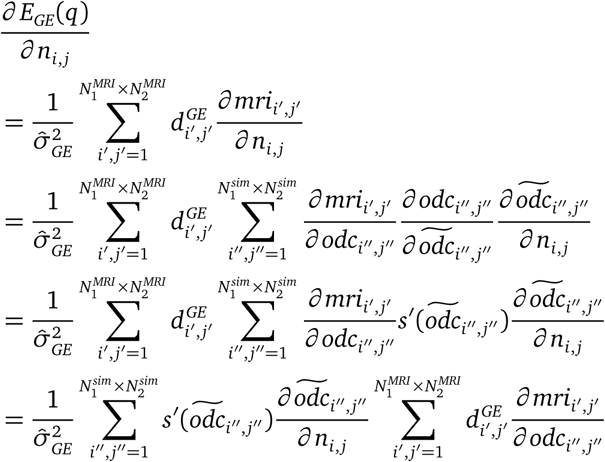

For both sums we apply D.8 (see D.2 for the definition of the zero-padding operation 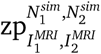):

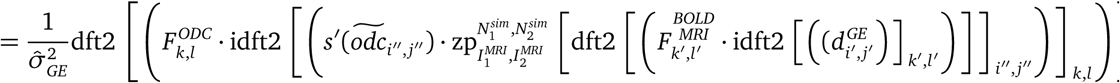

Similarily for the *SE* energy component we get:

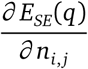

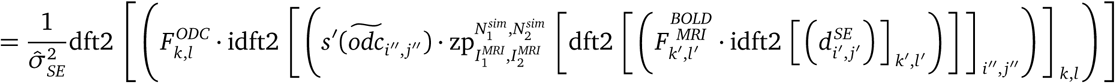

The contribution of the prior energy component is:

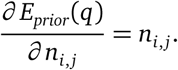

### Energy gradient with respect to ODC filter parameters

The derivatives of the *GE* energy component are:

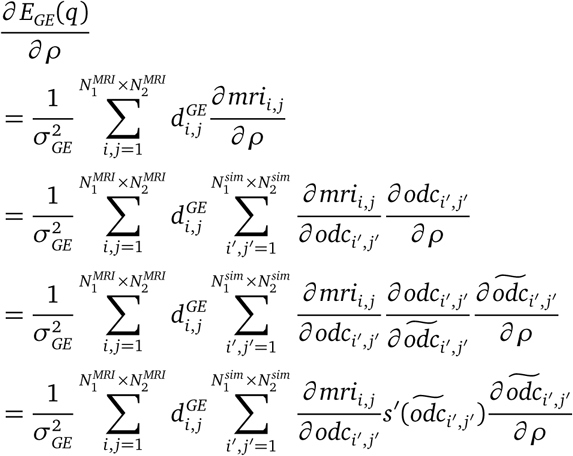

The last two factors together can be regarded as a pattern indexed by *i*′, *j*′. We apply D.7:

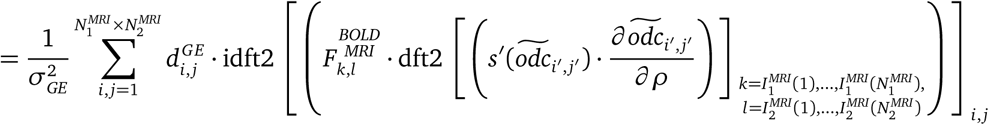

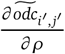 is the derivative of the ODC filter output 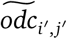 with respect to a filter parameter. We apply D.9:

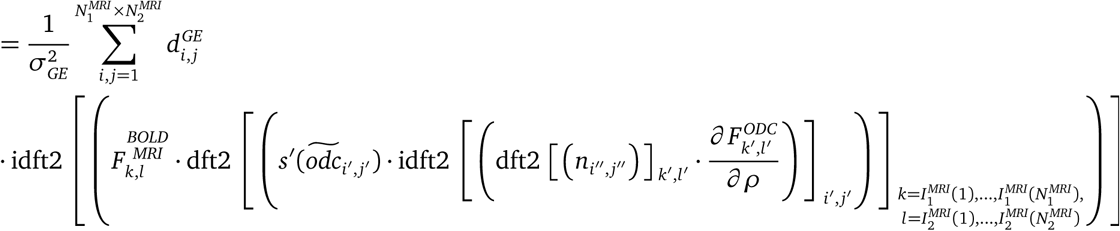

The derivatives of *E_GE_*(*q*) with respect to the remaining ODC filter parameters *δ*, *∈* and *θ* differ only in the derivative of 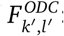 :

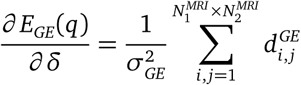

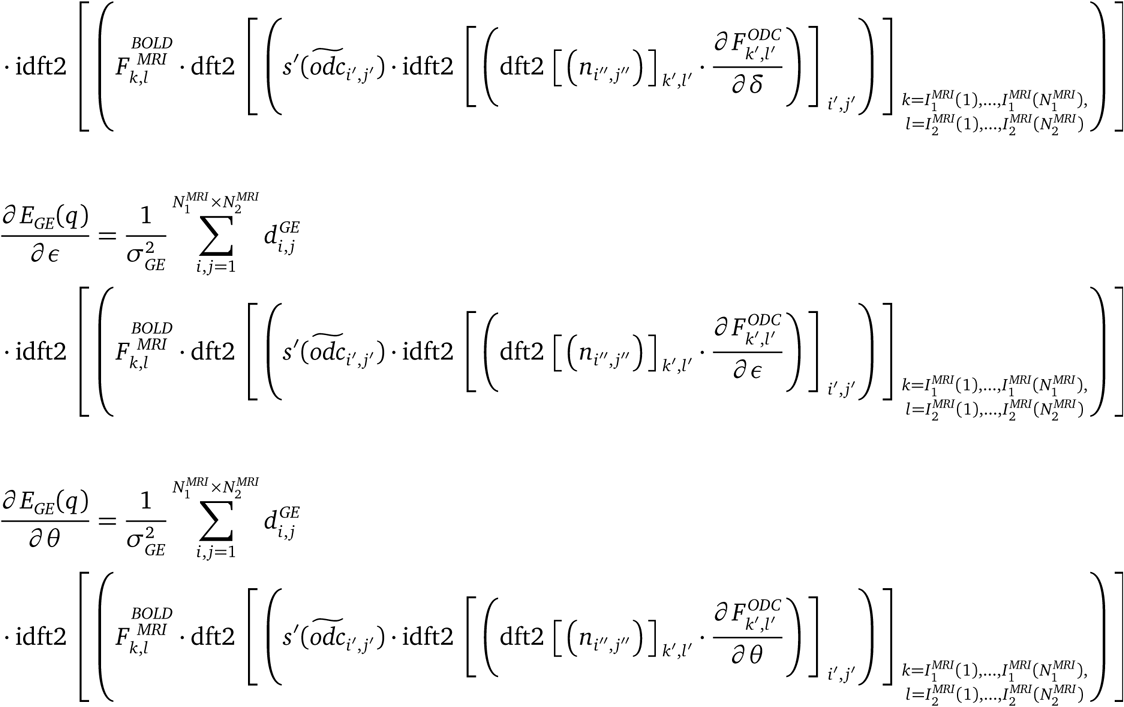

Similarily for the *SE* energy components we get:

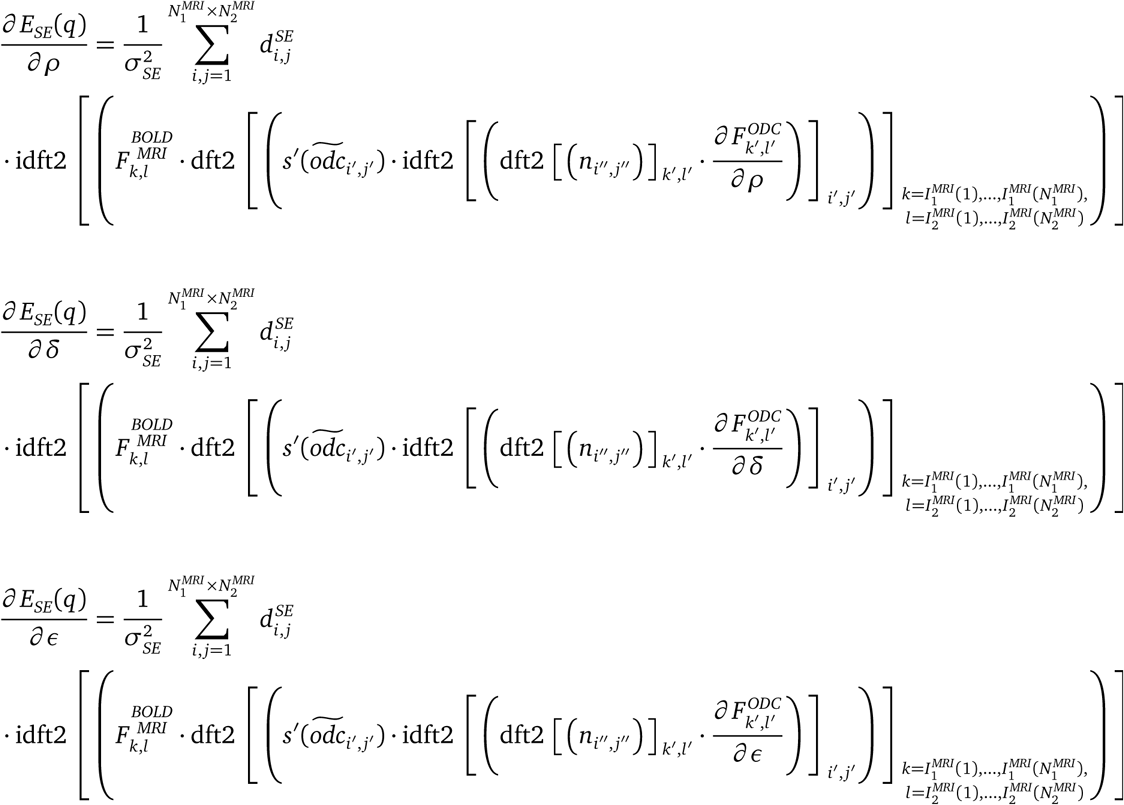

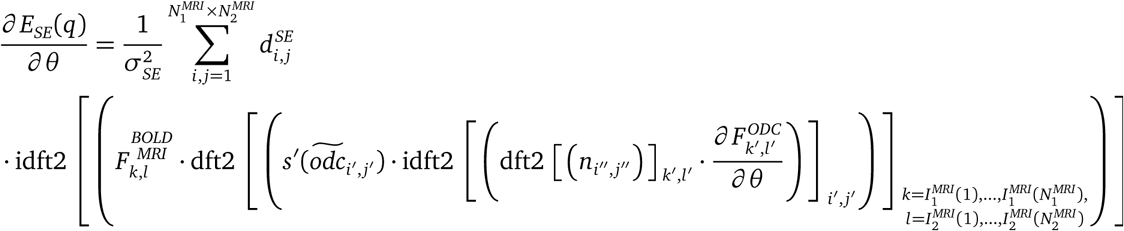

The contributions of the prior energy components are:

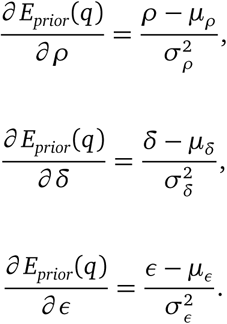

### Energy gradient with respect to the smoothness parameter

The derivatives of the GE component are:

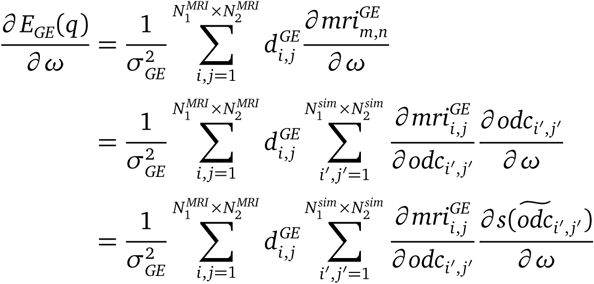

for the second sum we apply D.7:

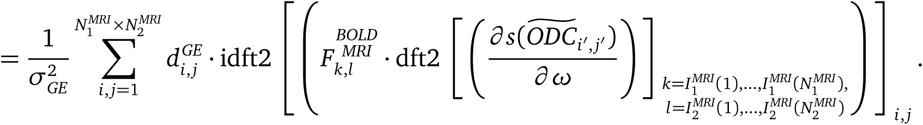

Similarily for the *SE* components we get:

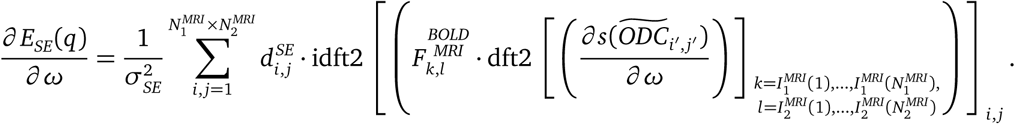

### Energy gradient with respect to BOLD PSF width

The derivatives of the GE component are:

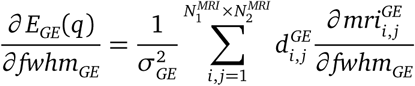

We apply D.9:

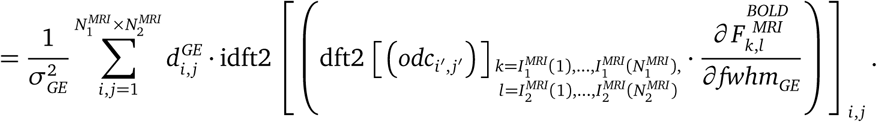

Similarily for the *SE* components we get:

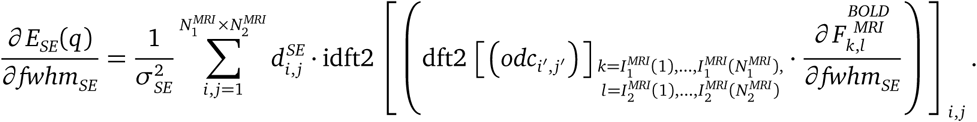

## Appendix D. Efficient computation of some expressions consisting of derivatives of filtering operations

Let us assume we have a pattern (*x_i, j_*) of size 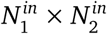 (the input), whose discrete Fourier transform is (*X_k,l_*). We filter (or equivalently: convolve) that pattern using a linear filter (*F_k,l_*) defined in frequency space and of size 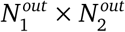. The filtering is carried out as:

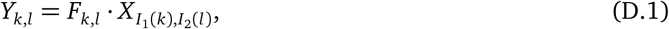

where (*Y_k,l_*) is the discrete Fourier transform of the filtered, 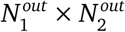 sized pattern (*y_m_,_n_*) (the output). *I*_1_ and *I*_2_ are index functions that allow to assign a specific subset of elements of (*X_k,l_*) to (*Y_k,l_*) (e.g. when restricting the spatial frequency space representation in order to simulate MRI sampling). Note that for 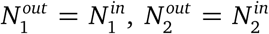, *I*_1_(*k*) = *k* and *I*_2_(*l*) = *l* this formalism describes a normal filtering operation in which the size and structure of the spatial frequency space is not altered.

Our goal here is to derive alternate formulations for some derivative expressions that result in a more efficient computation of gradients for models that contain such filtering operations.

### Definitions

We define a zero-padding operation 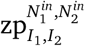 that allows us to up-sample patterns with size equal to the output pattern to match the size of the input pattern:

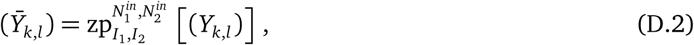

where 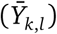 is of size 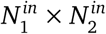 such that:

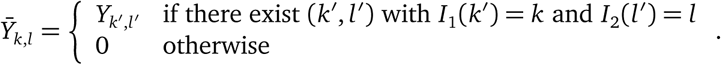

We use the following definitions of the two-dimensional discrete Fourier transform (*X_k,l_*) = dft2 [(*x_i_* and its inverse (*x_i, j_*) = idft2 [(*X_k,l_*)], where (*x_i, j_*) and (*X_k,l_*) are of size *N*_1_ × *N*_2_:

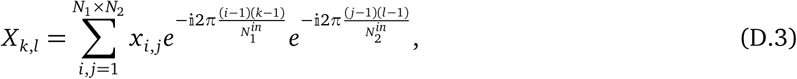

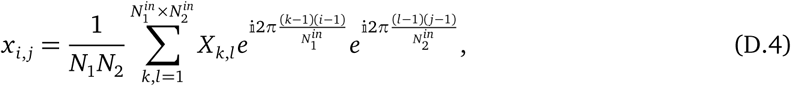

with partial derivatives given as:

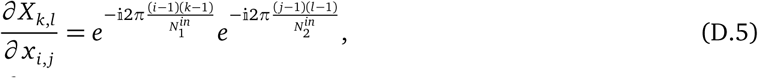

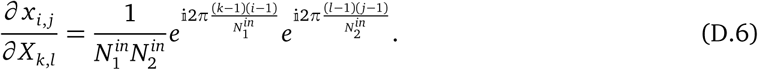

### Weighted sums of derivatives with respect to input

The first expression of interest is the weighted sum (weights given as (*g_i, j_*)) of the derivatives of an arbitrary output element *y_i, j_* with respect to all input elements *x_i′,j′_*. We derive a formula that allows to compute this expressions for all output elements *y_i, j_* simultaneously using one dft2 and one idft2 operation.

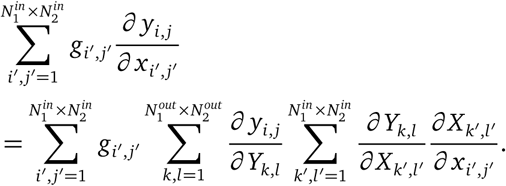

Because of D.1, the second to last term is equal to *F_k,l_*, if (*k′*, *l*′) = (*I*_1_(*k*),*I*_2_(*l*)) and 0 otherwise:

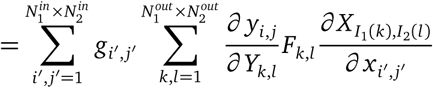

Furthermore making use of D.5 and D.6 we get:

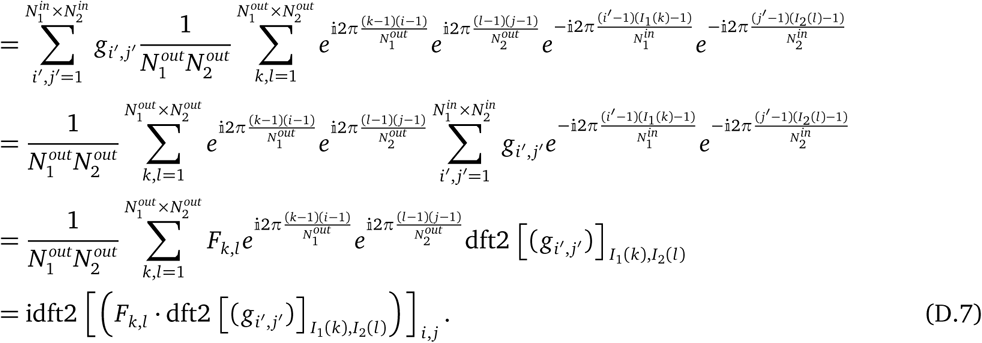

A related expression is the weighted sum of the derivatives of all output elements *y_i′,j′_* with respect to an arbitrary but specific input element *x_i, j_*. Again, we derive a formula that allows to compute this expressions for all input elements *x_i, j_* simultaneously using one dft2 and one idft2 operation.

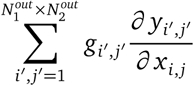

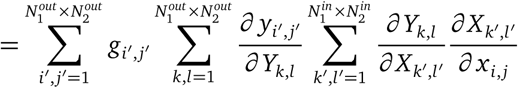

Because of D.1, the second to last term is equal to *F_k,l_*, if (*k′*, *l*′) = (*I*_1_(*k*), *I*_2_(*l*)) and 0 otherwise:

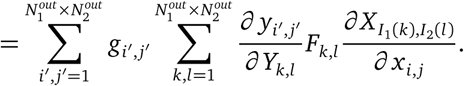

Furthermore making use of D.5 and D.6 we get:

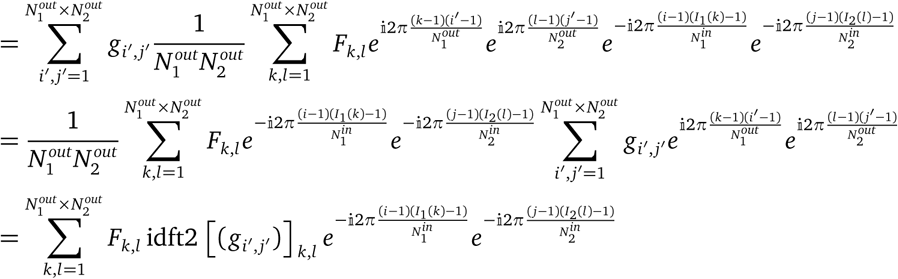

Instead of summing over the output indices 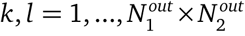 we can zero-pad *F_k,l_* idft2 [(*g_i′,j′_*)]_*k,l*_ and sum over the input indices 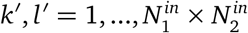. The additional elements are 0 due to the zero-padding and will not contribute to the sum.

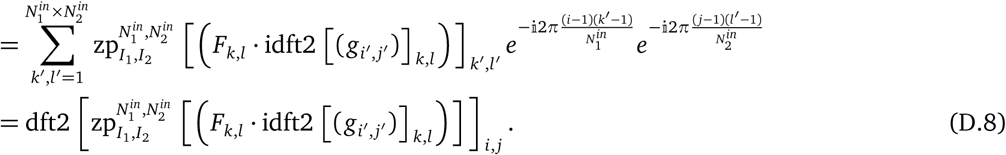

### Derivative with respect to a filter parameter

Let us assume that the filter representation (*F*_k,l_) depends on some parameter *q*. We derive a formula that allows to calculate the partial derivatives of all output elements *y_m_,_n_* with respect to *q* simultaneously using one dft2 and one idft2 operation.

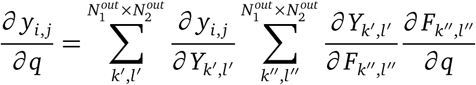

Because of D.1, the second to last term is equal to *X_I_*_1_(_k_′),_*I*2_(_l_′) if (*k*″, *l*″) = (*I*_1_(*k*′), *I*_2_(*l*′)) and 0 otherwise:

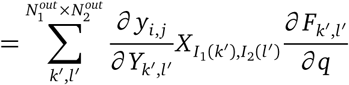

We apply D.6:

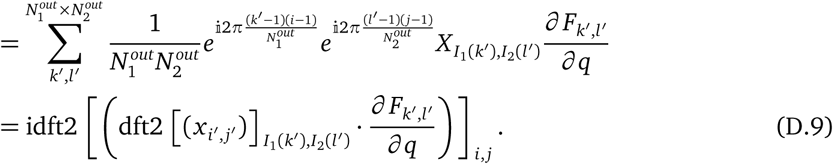

---

**Sup. Fig. 1.**
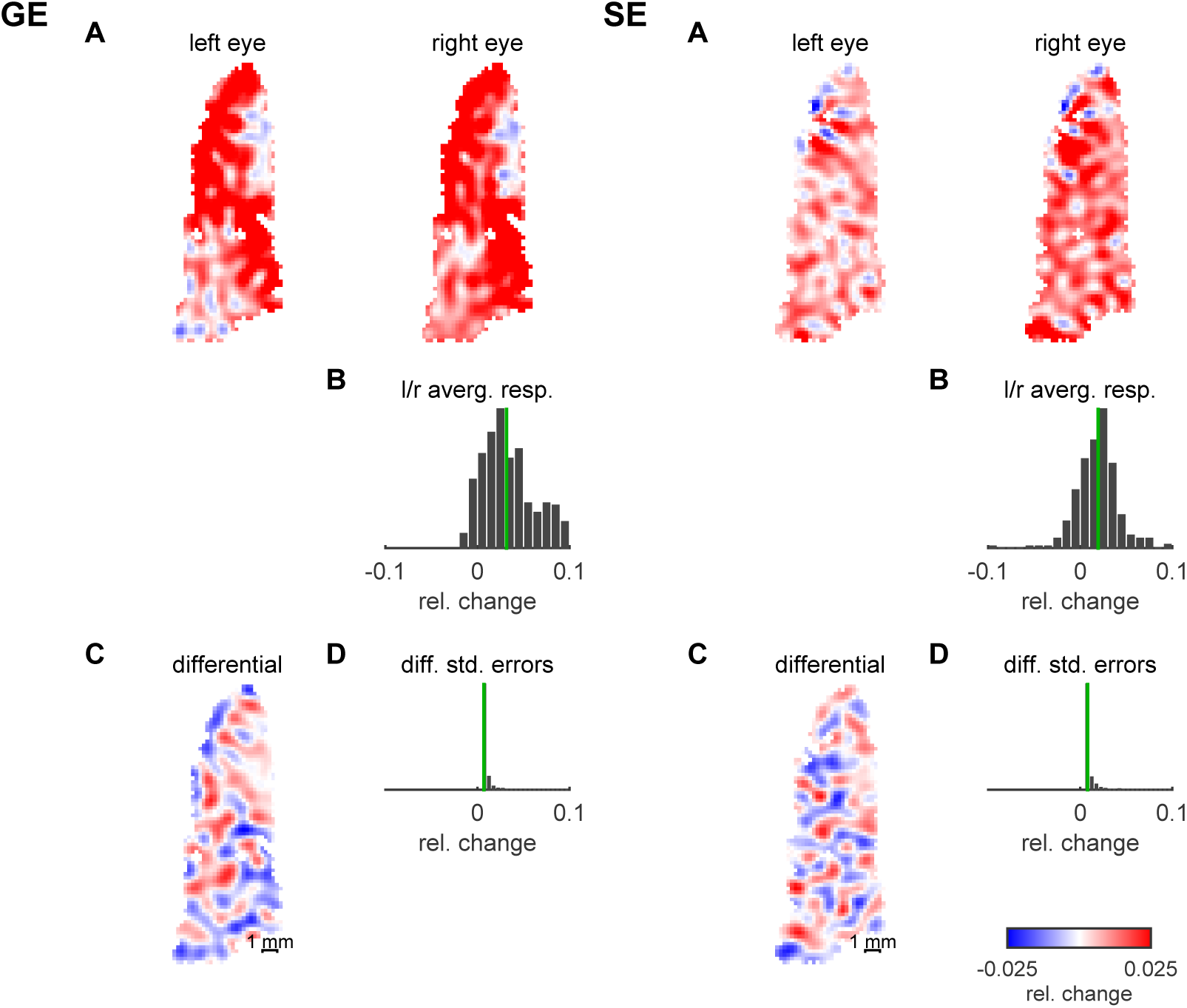
fMRI ODC data. Results from the GLM analysis of fMRI data from subject 2 for GE (left) and SE (right). **A** Responses to left and right eye stimulation relative to baseline. **B** The response maps to the left and right eyes from A were averaged. B shows the distribution of the average response. Its median (in green) was used to set the overall amplitude of the BOLD response model. **C** The difference between left and right eye responses yields the differential ODC map. **D** The distribution of standard errors of all differential responses. From this distribution we estimated the noise level used by the model. The color look-up-table applies to all response maps.

**Sup. Fig. 2.**
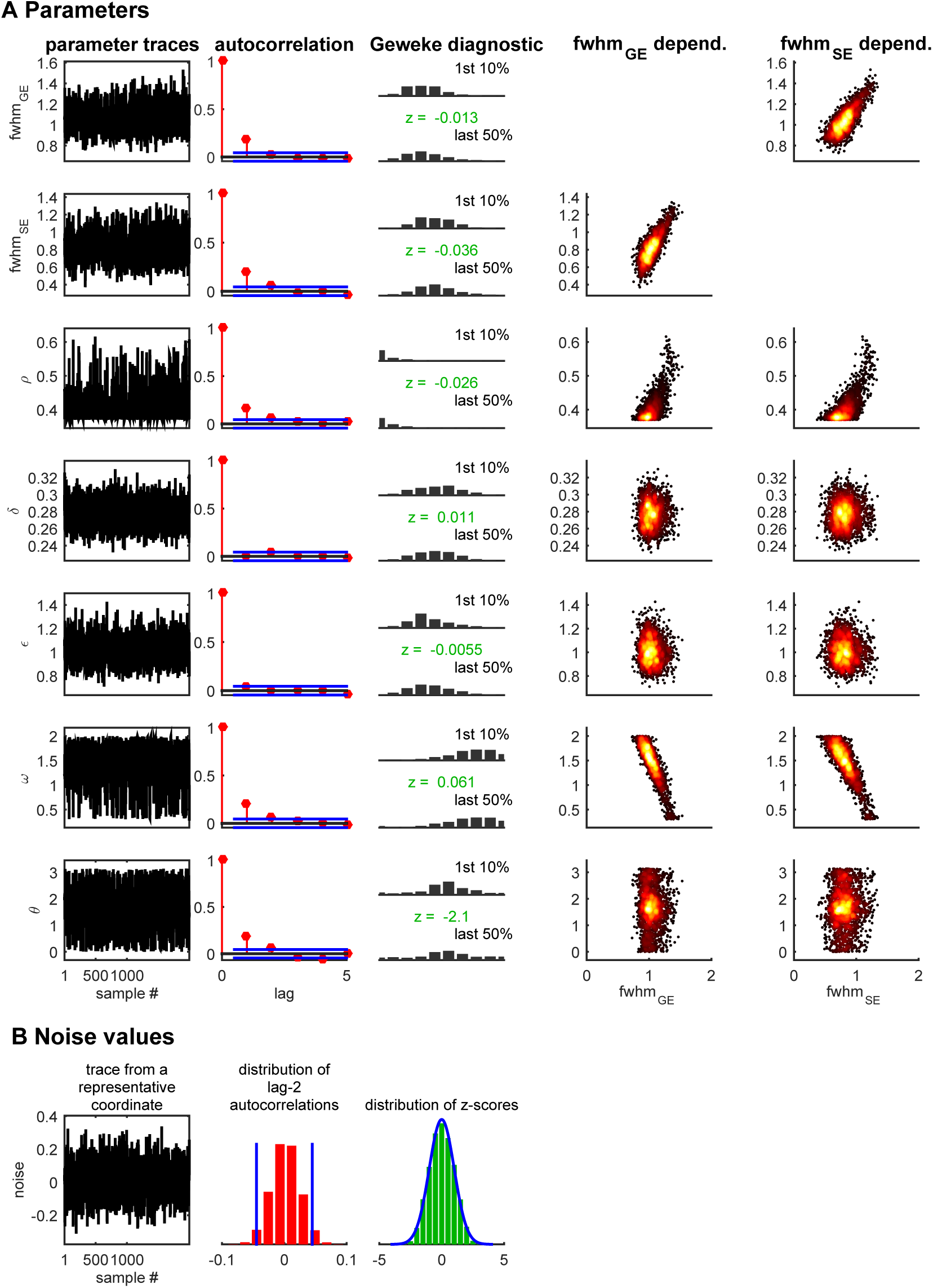
Convergence diagnostics of Markov Chain Monte Carlo sampling. Markov Chain Monte Carlo needs to run for a sufficient number of iterations in order to yield samples from the modeled probability distribution. Indications for convergence are: (1) stationarity of the parameter sampling distributions, and (2) sample autocorrelations decrease rapidly with increasing lag, relative to the total number of samples. This figure examines convergence for subject 2. The upper part of the figure (A) shows diagnostics for the standard model parameters. The bottom part (B) shows diagnostics for the white noise values that act as parameters to determine the ODC pattern. The first column (A and B) shows traces of the sampled parameters. For the noise values (B), one exemplary trace is shown from the center of the map. The second column (A and B) shows sample autocorrelations as a function of lag. The horizontal blue lines (A) indicate the 95%-confidence bounds around 0 for a white noise process. Consecutive samples (lag=1) show low autocorrelation. However, samples of lag 2 (and higher) show autocorrelation estimates that are comparable to those obtained from uncorrelated white noise. In B, the autocorrelation from the noise samples are summarized by the histogram of lag-2 autocorrelations from all coordinates together. Here, 95%-confidence bounds for a white noise process are indicated by vertical blue lines. The third column (A and B) presents the Geweke convergence (stationarity) diagnostic, which is a z-test (z-scores shown in green) for testing whether the means of the first 10% and last 50% of samples are different. In A, 2 histograms per each parameter show how similar their respective distributions are. In B, the z-scores from the noise samples are shown as a histogram together with a blue plot of the standard normal probability density representing the null-hypothesis of z=0. The last two columns (A) show the sample covariation between each parameter (vertical axis) and the GE and SE point-spread function FWHM (horizontal axis).

